# A synaptic-astrocytic proteomic signature associated with synaptopathy in Alzheimer’s Disease

**DOI:** 10.1101/2025.01.23.634408

**Authors:** Jessica Griffiths, Eléonore Schneegans, Harry Whitwell, Zhen Qiu, Beverley Notman, Dorcas Cheung, Nanet Willumsen, Paul M Matthews, Seth G N Grant, Johanna S Jackson

## Abstract

Synapse loss is the greatest correlate of cognitive impairment in Alzheimer’s Disease (AD) and offers a therapeutic avenue alongside disease-modifying therapies. However, the events preceding synapse loss in the human condition have not been well characterised. In this study, we describe a pseudotemporal profile of alterations in the synaptic proteome prior to excitatory synapse loss in human *post-mortem* brain AD tissue using synapse proteomics and synaptome mapping techniques. In a region with early-stage disease pathology, the most predominant changes were pre-synaptic and featured changes in metabolism and exocytosis. In a mid-stage disease state, alongside initial synapse loss, there was a dominance of inhibitory synaptic changes. In a region with late-stage disease pathology and profound synapse loss, post-synaptic changes were most prevalent with a range of canonical synaptic transmission pathways reduced and differential excitatory synapse subtype pathology. Synapse loss was associated with changes in astrocytic proteins which were enriched for those at peri-synaptic astrocytic processes, including an upregulation of complement activation and endocytosis; a signature that differed from the astrocyte cytosolic proteome. Taken together, this provides evidence of a cascade of events leading to synapse loss with multiple points for therapeutic intervention to alleviate cognitive decline in AD. Data are available via ProteomeXchange with identifier PXD056052.

## Introduction

Alzheimer’s Disease (AD), the most common form of dementia, represents a global health crisis, affecting over 57 million individuals worldwide and imposing significant emotional and economic burdens on families and healthcare systems^1^. With an estimated 153 million cases projected in 2050, there is a pressing need for research to facilitate the development of targeted preventative and disease-modifying treatments. Synapse dysfunction and loss are recognised as some of the earliest hallmarks of AD and associated with the development and progression of both behavioural and physiological disease characteristics^2,3^. Consequently, the synapse is considered a critical substrate linking cognitive symptoms and proteinopathy in AD, making it a promising target for novel therapies^4,5^.

Previous bulk^6,7^ and synapse proteomic human brain studies^8–13^ have provided valuable insight into the molecular underpinnings of AD, highlighting dysregulated synaptic proteins involved in signal transduction, vesicular assembly, metabolism, axonal transport, protein degradation, and neuroimmune interactions^14^. However, despite the plethora of synaptic changes identified, a comprehensive understanding of the association of the specific alterations that contribute to selective synapse vulnerability with the spatiotemporal progression of AD pathology^15^ is incomplete. Human *post-mortem* brain histopathological studies have demonstrated region-specific synapse loss^2,16–20^, synaptic protein loss^21–28^, and distinct morphological changes^18,29–33^, pointing towards selective vulnerability of specific molecular types and subtypes of synapses, but these have not been defined^34^. Likewise, a selective vulnerability of pre- vs post-synaptic components^35,36^ in preclinical AD models and of excitatory and inhibitory synapses^22,26^ in human AD have been shown although the molecular events responsible for these changes have not been characterised. To address these gaps, our study sought to investigate the progressive molecular changes at the synapse proteome and address its association with the loss of vulnerable synapses. For this, we used the progressive spatial nature of phospho-tau pathology in AD as a proxy measure of time course, where medial regions are affected early, followed by frontal regions and lastly the visual cortex^37–39^.

Sub-cellular compartmentalisation serves to optimise neuronal function^40^ however, intrinsic (cell autonomous) processes such as cellular transport^41^ and non-cell autonomous processes (affected by other cells), such as glial cell interactions^42^ influence the health and function of compartments such as the synapse. Biochemical fractionation of neuronal compartments can be obtained and subjected to proteomic mass spectrometry. This method has previously been used to evaluate the influence of each cellular compartment on synapse function in AD^43^. Furthermore, using the SynPER preparation method used here, non-neuronal synaptic components can be retained^44^ such as peri-synaptic astrocytic processes (PAPs), crucial for synapse homeostasis and memory expression^42^, and allows the interrogation of their proteome in AD. We therefore took advantage of the SynPER preparation to characterise the astrocytic proteomic changes in paired analyses alongside the synaptic fraction to distinguish extrinsic influences from synapse-autonomous processes.

We utilised synapse proteomics and synaptome mapping techniques to define a regional profile of synaptic protein alterations and analyse NMDA receptor subunit GluN1-positive excitatory synapse loss in late-stage AD and non-diseased control *post-mortem* samples. Our findings provide a detailed characterization of regional molecular pathology at the synapse and reveal differential excitatory synapse subtype vulnerability in AD. We performed an integrative analysis of localised synaptic and associated cellular differences across the course of AD pathology to derive a disease pseudotime trajectory, and discovered an astrocytic signature associated with the observed differential synapse subtype loss, pointing towards a key role for these glial cells in AD pathology.

## Results

Twenty-six *post-mortem* brains were assessed using synaptomic approaches (***Supplementary Fig 1***), 13 of which had been neuropathologically diagnosed with AD (Braak stage V-VI) and 13 were non-diseased control (Braak stage 0-II) cases (***Supplementary Data 1***). Three cortical brain regions were selected to represent regions of varying degrees of AD pathology: the visual cortex (VC, Brodmann area 17), prefrontal cortex (PFC), and middle temporal gyrus (MTG). Due to tissue availability, a subset of cases and regions were used for each experimental procedure (***Supplementary Data 1***).

### Selective vulnerability of synaptic components at different stages of AD

In the synaptic fraction, we identified over 6000 proteins across all brain regions with 5688 ±142 (1 standard deviation) proteins per sample. Approximately 60% of proteins exhibited a coefficient of variation (CV) <30% within distinct brain regions (***Supplementary Fig. 11a-d***). In the cytosolic fraction, we identified fewer proteins compared to the synaptic fraction with a lower proportion of proteins with a CV <30%, indicative of the diversity of cell-types within the fraction (***Supplementary Fig. 12a-d***).

To ensure enrichment for synaptic proteins in the synaptic fraction, we performed several validation steps (***Supplementary Fig 2a-c***). First, we compared the synaptic fraction against a consensus list of proteins related to individual subcellular compartments, as previously described^12^. This analysis revealed a significant enrichment of proteins localised to the postsynapse (3.3-fold, p.adjusted = 3.12e^-225^) and presynapse (1.6-fold, p.adjusted = 2.64e^-05^) (***Supplementary Fig 2a***). We also observed a lower abundance of specific pre- and post-synaptic proteins in the cytosolic fraction compared to the synaptic fraction (***Supplementary Fig 2b***). Notably, 77% of the proteins in the synaptic fraction overlapped (***Supplementary Fig 2c***) with those reported in two recent synapse proteomic studies^12,45^. Lastly, for the initial assessment of the synaptic proteome without cytosolic proteome integration, we used a bioinformatic filter with an inclusion list of 5667 synaptic proteins (***Supplementary Data 5***^12^), identifying 3333 high-confidence synaptic proteins across all regions.

Having validated our sample preparation protocols, we next asked how many proteins were differentially expressed (DEPs) between AD and controls in the three brain regions. Differential expression analysis of these 3333 proteins revealed differences in the VC (412 DEPs: 6.9% increased, 5.4% decreased), PFC (380 DEPs: 6.6% decreased, 4.8% increased), and MTG (504 DEPs: 7.8% decreased, 7.3% increased) in AD compared to control brains (***Fig 1a, Supplementary Data 6***). Euclidean clustering based on DEP relative abundance demonstrated a clear separation of the AD and control groups in all three regions (***Supplementary Fig 3***). Overall, we reported 656 DEPs that were not identified in two whole-tissue proteomic AD studies^6,7^ and a recent synapse proteomic AD study^12^ (***Supplementary Fig 4a***, ***Supplementary Data 7***). In all three regions, several canonical synaptic proteins were differential expressed (e.g. BSN, NRXN3, SYT12), while brain region-dependent changes were also noted. For instance, GABA_a_ receptor subunits were downregulated in the PFC (GABRA1, GABRA2, GABARG2, GABRB3) but not in the VC or MTG. Isoform-specific changes were also observed, with membrane-trafficking synaptotagmin family members either being downregulated in all regions (SYT12), downregulated in a specific region (SYT1, SYT12 and SYT17 in the MTG), or upregulated in a specific region (SYT11 and SYT5 in the VC).

**Figure 1:**
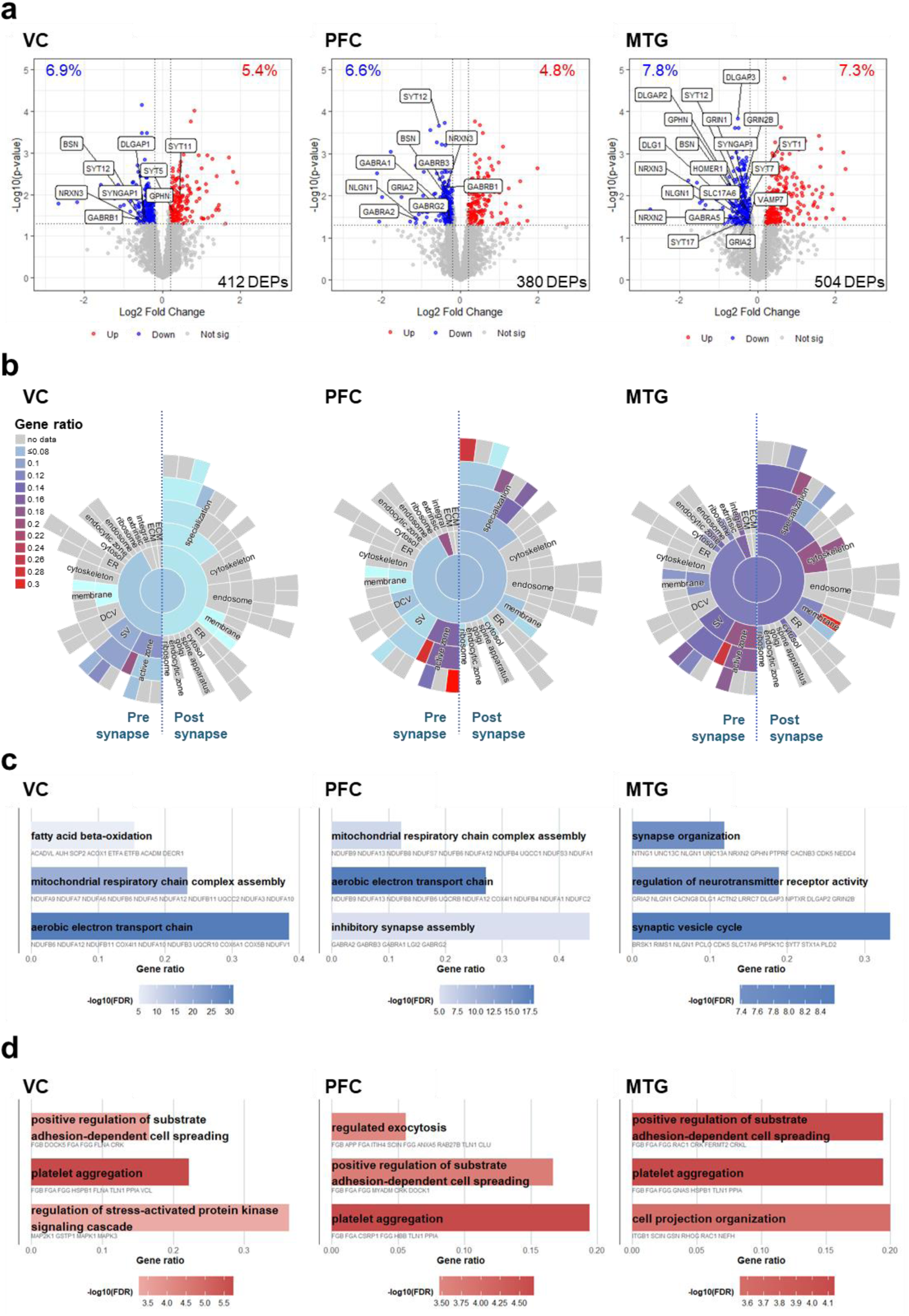
Differential synaptic protein expression across cortical regions with Alzheimer’s Disease. **(a)** Volcano plot depicting differentially expressed proteins (DEPs) between AD and control groups in the VC, PFC, and MTG. Canonical synaptic proteins of interest are highlighted. **(b)** Sunburst plots displaying the gene ratio calculated as the ratio of significant genes versus all annotated genes per SynGO cellular component term in the VC, PFC, and MTG (one-sided over-representation Fisher’s exact test with multiple testing correction using FDR). **(c,d)** Bar plots illustrating the relative functional enrichment against GO biological process terms on DEPs that have a negative **(c)** and **(d)** positive fold change in the VC, PFC, and MTG in AD relative to control (colour, log_10_(FDR), one-sided over-representation Fisher’s exact test).

We next compared the abundance of proteins in subsynaptic compartments through synaptic enrichment analysis using the annotations within the SynGO database^46^. A significant representation of DEPs was observed in the synapse compartment in all cortical regions (***Fig 1b; Supplementary Data 8***). In the VC, we noted differential enrichment between the presynaptic and postsynaptic compartments, with the most significant enrichment in the presynaptic active zone, particularly in cytoplasmic components. In the PFC, pronounced presynaptic dysfunction affected both cytoplasmic and membrane components of the active zone, along with dysfunction in the postsynaptic density, especially in its anchored component. This pathology was more widespread in the MTG, where enrichment spanned multiple pre- and post-synaptic compartments, indicating broader synaptic dysfunction. Additionally, we examined potential differences between excitatory and inhibitory synapse classes. Using a consensus list of excitatory and inhibitory synaptic proteins (***Supplementary Data 9***^22^), we found that proteins unique to both classes were differentially expressed in all regions of AD (***Supplementary Fig 5***). However, a greater percentage of inhibitory DEPs were specific to the PFC, while a greater percentage of excitatory synaptic proteins were differentially expressed in the MTG compared to the PFC and VC (***Supplementary Fig 5a-d***). These findings suggest a region-dependent imbalance in excitatory/inhibitory synaptic function.

Gene ontology (GO) pathway enrichment analysis was employed to identify biological processes associated with proteins that had a negative (***Fig 1c***) or positive (***Fig 1d***) fold change in AD compared to that of the control group (***Supplementary Data 10***). While some biological pathways were shared across regions, others represented brain region-dependent changes. For example, proteins that were downregulated were significantly enriched in mitochondrial-related pathways in the PFC and VC, indicating synaptic metabolic dysfunction in these brain regions^47^. Additionally, we observed a reduction in pathways related to inhibitory synapse assembly and transmission in the PFC, although not in the VC where inhibitory synapse loss has been previously reported^26^. In the MTG, we reported an enrichment in pathways associated with synaptic organisation and function, confirming overt synaptic dysfunction in the MTG.

Pathways enriched with proteins exhibiting a positive fold change were conversely shared across regions suggesting pathological processes common to all. These included pathways related to the regulation of substrate adhesion, cell-spreading and platelet aggregation^48,49^. We further examined the relationship between these proteins and AD risk genes. Utilizing MAGMA, a geneset analysis tool for GWAS data, we found these DEPs with increased relative abundance were enriched in risk genes across all regions, including clusterin, the gene of which has been linked to late-onset AD^50–52^ (***Supplementary Fig 4b***). Taken together, these data suggest region-specific synaptic profiles in AD including differential vulnerability of pre- and post-synaptic subcompartments, excitatory and inhibitory imbalances, and overall synaptic changes as pathology progresses.

### Brain-region-independent synapse proteomic signatures associated with disease progression

We next interrogated synaptic proteins differentially expressed between AD and control groups in all regions to identify general disease-associated synaptic differences that could suggest potential soluble biomarkers. While we observed a greater number of proteins altered in a brain region-dependent manner, 70 proteins showed brain-region-independent alterations in AD, with 30 being downregulated (***Fig 2a***) and 40 upregulated (***Fig 2b***).

**Figure 2:**
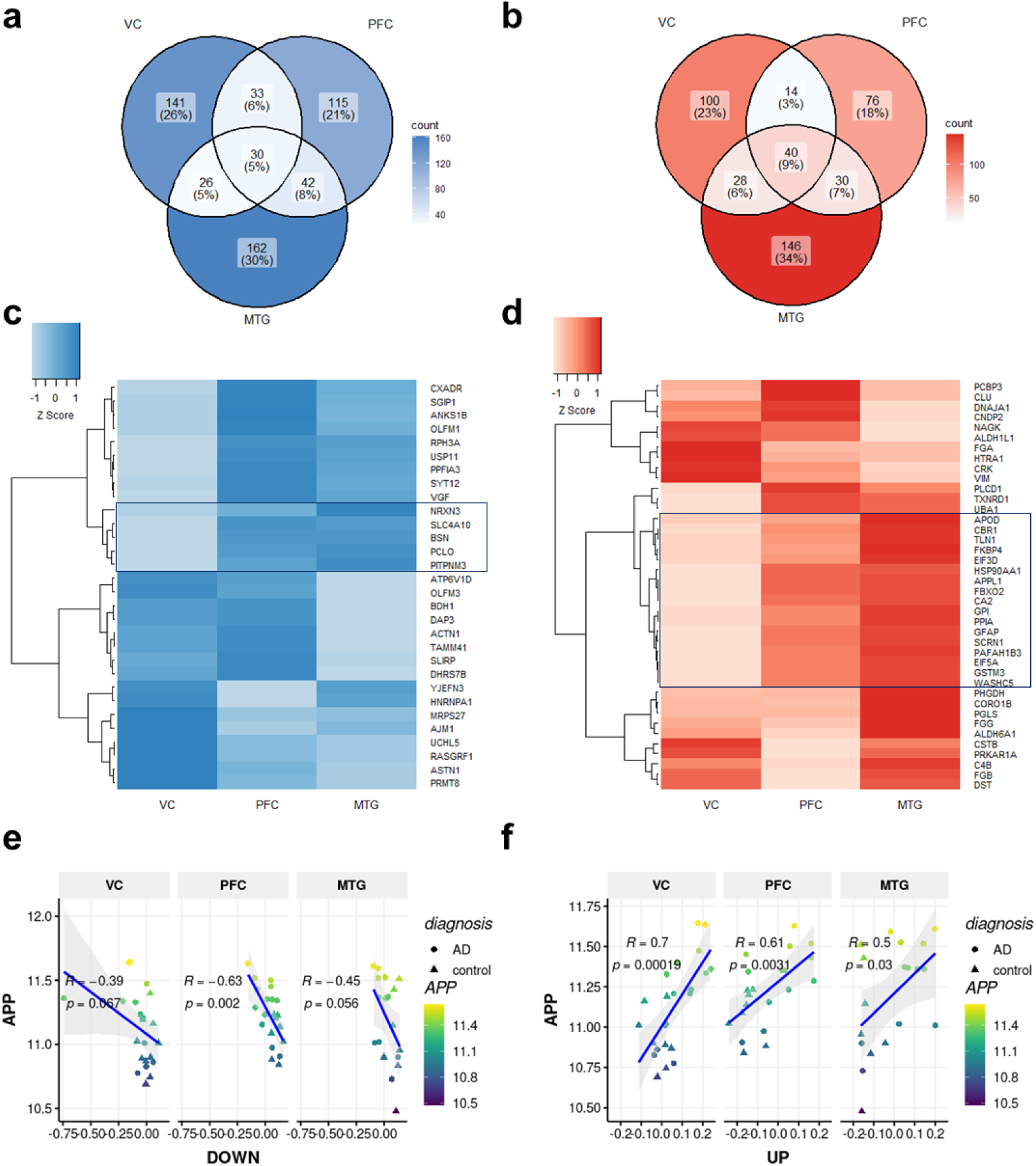
Brain region dependent and independent synaptic signatures with AD. **(a,b)** Venn diagrams illustrating the overlap of DEPs that are either downregulated (**a**) or upregulated (**b**) in the VC, PFC, and MTG in AD compared to control groups. DEPs are filtered based on a log_2_ fold change threshold >0.2 and significance level of p<0.05. The percentage of proteins in each intersection is rounded to the nearest whole number. **(c,d)** Heatmaps displaying normalised log_2_ fold change (row Z-score) of brain region-independent DEPs that are **(c)** downregulated and **(d)** upregulated in all three regions. Euclidean clustering based on the fold change identifies proteins showing stepwise increases in differential expression from the VC to the PFC to the MTG (highlighted in black boxes). **(e,f)** Linear regression analysis of eigenprotein expression values derived from brain region-independent DEPs identified in **(c)** and **(d)** (black boxes), in relation to APP protein abundance, stratified by brain region.

Among the proteins upregulated were astrocyte-associated proteins such as CLU, ALDH1L1, GFAP, and VIM, and complement refinement protein, C4B. Of note, the astrocytic protein MFGE8 was upregulated in the PFC but not differentially expressed in the other regions. Downregulated proteins included postsynaptic scaffolding proteins such as BSN and PCLO, proteins involved in receptor motility like RPH3A and NRXN3, and VGF, a dysregulated protein in neurodegenerative and psychiatric disorders that has been proposed previously as a therapeutic target^53^.

Clustering based on relative expression of differentially expressed proteins common to all regions revealed groups of proteins exhibiting stepwise increases in differential expression from the VC to the PFC to the MTG (***Fig 2c-d, black box***), suggesting progressive dysregulation as AD pathology advances. Indeed, the computed eigenprotein expression for both downregulated and upregulated proteins showed positive and inverse correlations respectively, with the abundance of amyloid precursor protein (APP) in the synaptic fractions (***Fig 2e-f***). Furthermore, this relationship was also reflected when assessing histopathological measures of percentage immunoreactive area of amyloid-beta plaques (Aβ; 4G8) and phosphorylated tau (pTau; PHF1) related to disease progression (***Supplementary Fig 6, Supplementary Data 11***). Taken together, these data highlight the progressive nature of synaptic changes in AD that are associated with increasing pathological load and may provide pharmacodynamic measures to be assessed in preclinical or cellular models during therapeutic development.

### Differential excitatory synapse subtype vulnerability in AD

To understand how changes in the synapse proteome may be associated with synapse loss and vulnerability, we next assessed synaptic protein expression using immunolabelling of tissue sections from AD and control cases. We focussed on the NMDA receptor subunit GluN1, the obligatory subunit of this ionotropic receptor which plays a key role in synaptic plasticity and learning^54^. GluN1+ synaptic puncta were immunolabelled, imaged, quantified and classified into 4 morphological subtypes (S1-S4) based on their punctum intensity and shape parameters (***Fig 3a-c***), as similarly conducted in the mouse brain^55,56^. The total GluN1+ puncta density (number per 100µm^2^) was quantified across the cortical grey matter in each brain region (***Fig 3c***). An unpaired t-test revealed no significant change in GluN1+ synapse puncta density in the VC; however, there was a significant decrease in the PFC and MTG in AD compared to control (***Fig 3c, Supplementary Data 12***). Next, we examined the GluN1+ subtypes and found differential pathology (***Fig 3c, Supplementary Data 12***). Strikingly, we found a significant loss of S2 and S4 excitatory synapse subtypes, an increase in the S3 subtypes, and no change in S1 in MTG and no changes in the PFC or VC which was associated with an overall increase in puncta size in the MTG (***Fig 3c***) Overall, this shows that there is not a loss of all NMDA receptor-expressing synapses in the MTG, but differential excitatory synapse subtype vulnerabilities.

**Figure 3:**
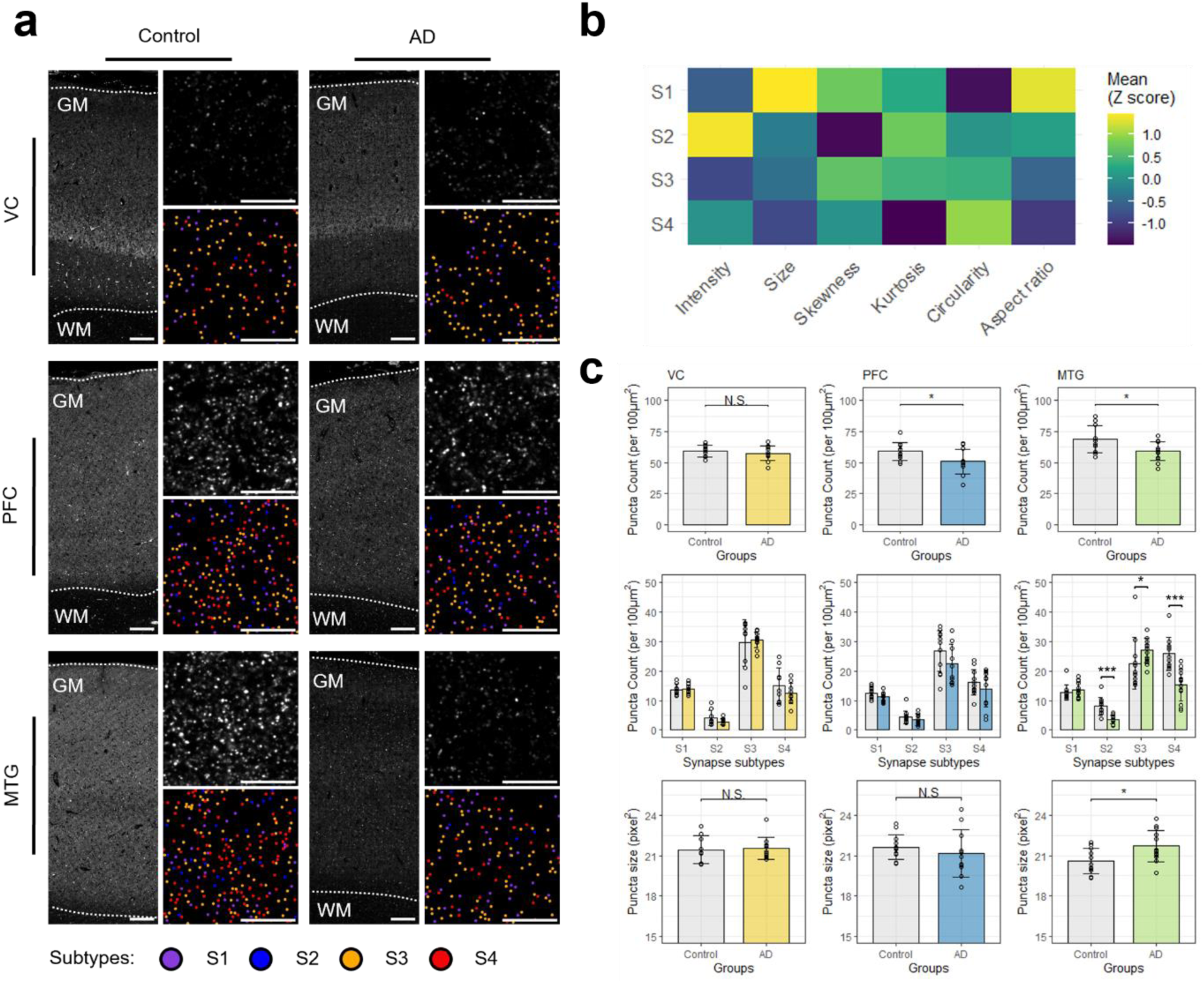
GluN1-expressing synapse subtypes are differentially affected by AD pathology in the MTG. **(a)** Representative immunohistochemical images showing GluN1+CY5+ synaptic puncta in three cortical brain regions in AD and control cases. High-magnification images taken from superficial layer II of the cortex with mapped GluN1+ synapse subtypes illustrated below (S1-S4) (GM, grey matter; WM, white matter; scale bar, 10µm). **(b)** Mean values of synaptome parameters (puncta intensity, size, skewness, kurtosis, circularity, aspect ratio) for GluN1+ subtypes. Values are normalised as Z-scores for each parameter. **(c)** Bar plots displaying mean densities of GluN1+ synaptic puncta (unpaired t-test, *p≤0.05), densities of GluN1+ synapse subtypes (multiple unpaired t-tests or Mann-Whitney U tests with Benjamini-Hochberg correction, *p≤0.05, **p≤0.01, ***p≤0.001) and GluN1+ synaptic puncta size (unpaired t-tests or Mann-Whitney U tests, *p≤0.05) in AD and control groups. Jittered dots represent individual case values.

### Synaptic-cytosolic proteomics integration identifies a discordant glia-synapse relationship

To further understand the mechanisms that co-ordinate cell-wide (synaptic and cytosolic) and local synaptic changes in relation to disease progression and synapse subtype pathology, we conducted an unsupervised integration of the synaptic and cytosolic proteomics using the Multi-Omics Factor Analysis (MOFA) model^57^ to identify shared axes of variation (i.e. *factors*) relevant to AD pathology. Using K=10 factors, the model explains up to ∼23% of the variation in the synaptic fraction and ∼41% in the cytosolic fraction (***Fig 4a, Supplementary Data 13)***. We defined the AD-correlated factors as 1,2,3, given the significant correlation between AD neuropathology variables and these factors (range of R^2^ value from 0.38 to 0.64, *p≤0.05,* ***Fig 4b***). These AD-correlated factors were used in a semi-supervised approach to derive a pseudotime measurement, representing disease progression, where a higher pseudotime represents a longer time spent in disease state. Briefly, this involved deriving a disease trajectory from the UMAP of AD-correlated factors, and to then rank samples according to this trajectory within early (0-II) and late (V-VI) Braak tiers (***Supplementary Fig 7***). In relation to the vulnerable GluN1+ synapse subtypes in AD, we observed that pseudotime was significantly negatively correlated with the summed density of S2 and S4 synapse subtypes in the MTG region (***Fig 4c***). In contrast, an opposite trend was found for the S3 subtypes. We next selected factor 3 for downstream analyses because its variation was derived from both synaptic and cytosolic fractions, unlike factor 1, which was primarily explained by the cytosolic fraction, and factor 2, which was mainly explained by the synaptic fraction. Specifically, factor 3 accounted for approximately 5% of the variance in the synaptic fraction and 3% in the cytosolic fraction. More so, factor 3 showed significant enrichment in SynGO pathways, indicative of synaptic function changes related to AD (***Supplementary Fig 8***). Constructing synaptic-cytosolic co-expression networks based on features in the AD-correlated factor 3 revealed three AD-correlated modules (***Fig 4d, Supplementary Data 14***). Two of these modules were prioritised for further analysis due to their strong associations with AD pathology and lack of age confound. Using EWCE^58^ to test for cell-type enrichment against a reference dataset from the Allen Human Brain Atlas^59^, we found that F3_DOWN_ME4 was related to neuronal cell types, whereas F3_UP_ME2 was enriched for glial cells, including astrocytes (***Fig 4d***). Linear regression analysis of module eigenvalues revealed an inverse relationship between glia and synapse signatures during AD progression (***Fig 4e***) further highlighting the role of astrocytes at the synapse.

**Figure 4.**
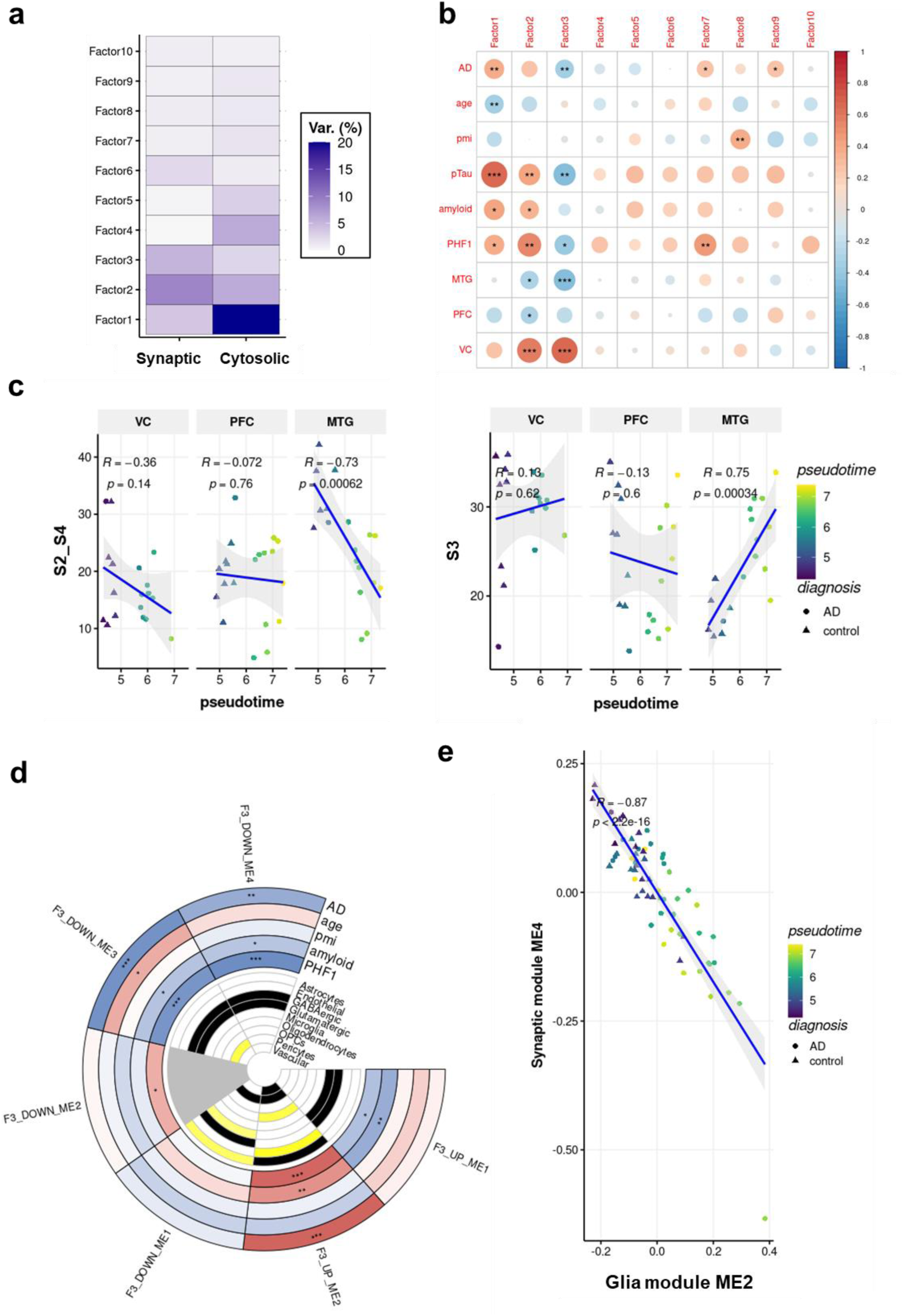
Synaptic-cytosolic proteomics integration unravels divergent glia-synapse dynamics. **(a)** Variance decomposition showing the percentage of explained variance per proteomics synaptic and cytosolic fractions and factor of the MOFA model with 10 factors. For each fraction, the heatmap shows the percentage of variance explained by the respective factor. **(b)** Heatmap showcasing the Pearson’s correlation matrix of clinical and pathological parameters with 10 MOFA factors derived from bulk synaptic-cytosolic integration. **(c)** Linear regression of the summed densities (count per 100µm^2^) of S2 and S4 synapse subtypes (S2_S4) and of S3 subtype against pseudotime, stratified by brain regions. **(d)** Circos plot visualising module (ME) trait relationships Pearson’s correlations indicating significant associations with key AD features (positive association in red, negative in blue). Cell type enrichment in the inner circle indicates significant enrichment in black (p≤0.05), near significance in yellow (p>0.05 – p<0.1), and non-significant in white. pmi = post-mortem interval **(e)** Linear regression of synaptic module eigenvalue as a function of the astrocyte module eigenvalue. Pearson’s correlation p-values derived from t-test (*p≤0.05, **p≤0.01, ***p≤0.001)

We further investigated the proteins in F3_DOWN_ME4, which primarily includes proteins from the synaptic fraction, such GluN1, (***Fig 5a***) and used in our synaptome mapping (***Fig 3***). Linear regression analysis of the summed density of S2 and S4 synapse subtypes as a function of the module eigenvalue revealed a downregulation of the module corresponding to the decrease in the summed S2 and S4 density, specifically in the MTG region. This relationship mirrored the abundance of the GluN1 protein and summed S2 and S4 density, underscoring the critical relationship of GluN1 and synapse subtypes (***Fig 5b***). Additionally, we observed significant enrichment of synaptic transmission pathways within this module, further validating its role (***Fig 5c***). This module indicates localised regulatory mechanisms or protein networks confined to the synaptic compartment, involving proteins essential for neurotransmitter release, synaptic receptor dynamics, and synaptic scaffolding, without counterparts or direct interactions with cytosolic proteins. The GluN1+ module highlights the presence of highly specialised synaptic processes that operate independently of broader cellular changes, emphasising the compartmentalised nature of synaptic regulation in the context of AD pathology.

**Figure 5.**
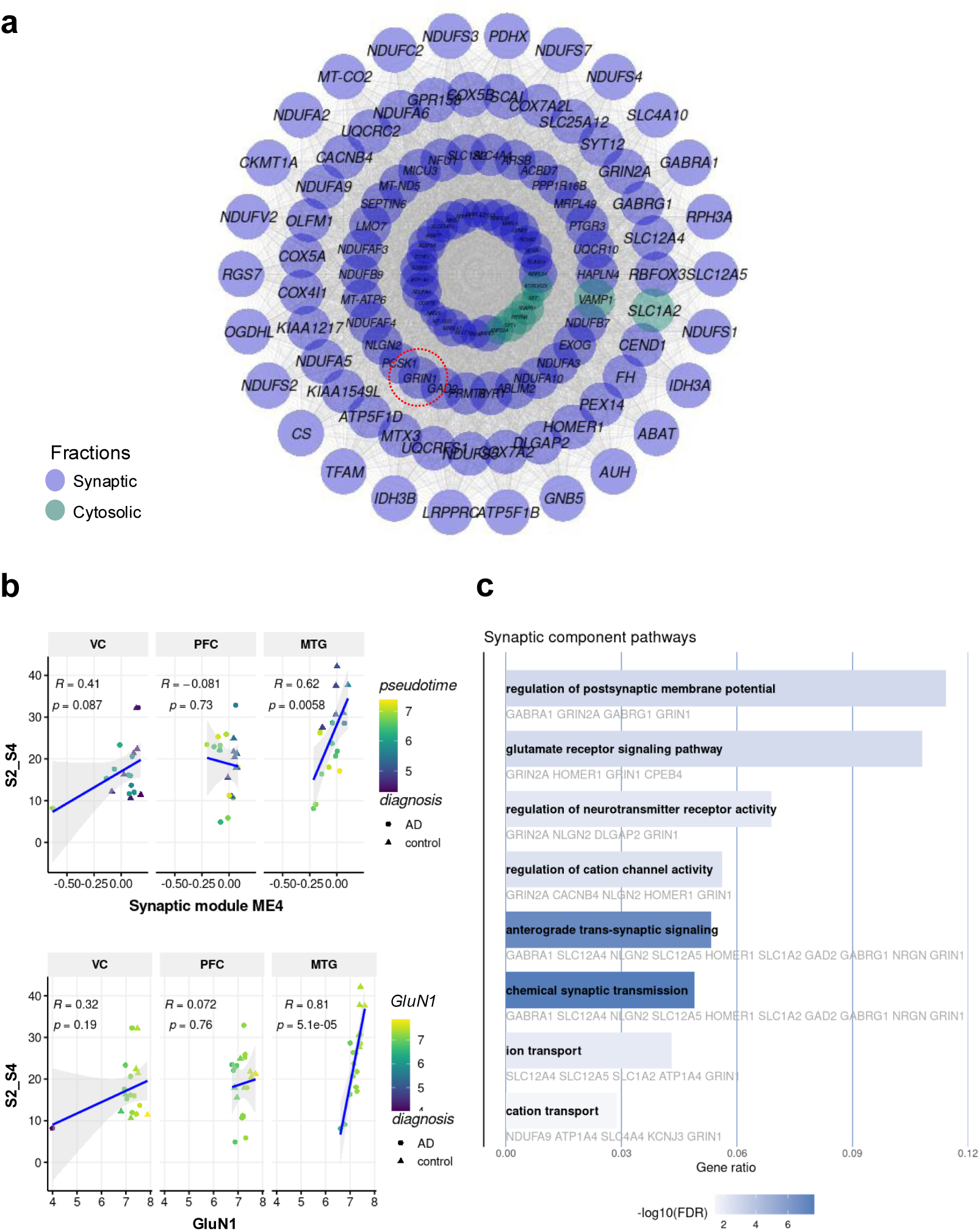
Summed S2 and S4 synapse subtype loss is associated with synaptic proteomic differences. **(a)** Schematic co-expression network diagram showing module membership of protein from the cytosolic fraction (green) and protein from the synaptic fraction (purple) in the GluN1+ (GRIN1; red circle) synaptic module. Node size is proportional to connectivity (node with a degree >50 shown on the graph for clarity), and nodes are separated into four quantiles representing each layer. **(b)** Linear regression of summed density of S2 and S4 synapse subtypes (S2_S4) as a function of the synaptic module ME4 eigenvalue (F3_DOWN_ME4) (top) and GluN1 protein abundance (bottom). **(c)** Bar plots of the functional enrichment analysis against GO biological process terms (containing GluN1) on the synaptic fraction proteins. Gene ratio represents the ratio of significant genes versus all annotated genes per pathway (colour: log_10_(FDR) one-sided over-representation Fisher’s exact test).

In contrast, our exploration of the F3_UP_ME2 glial module displayed proteins from both the synaptic and cytosolic fractions (***Fig 6a***). Linear regression analysis of the summed density of S2 and S4 synapse subtypes as a function of the glia module eigenvalue confirmed a significant relationship (***Fig 6b***). To determine if the astrocyte enrichment observed in the synaptic fraction could be attributed to proteomic changes associated with PAP/ASCs, we conducted cell-type enrichment analyses for both fractions with and without PAP/ASC proteins (***Fig 6c, Supplementary Data 15***^60,61^). Astrocyte enrichment in the synaptic fraction was no longer significant when PAP/ASC proteins were removed, unlike in the cytosolic fraction. This suggests the presence of other astrocytic components or functions within the cytosolic milieu, and that the astrocytic signature in the synaptic fraction is largely explained by changes in proteins localized at the PAPs. The synaptic fraction of this module, which consistently overlapped with the ASC junction module identified by Niu et al.^60^ (***Supplementary Fig 9***), showed enrichment in pathways related to the regulation of neuron death, receptor-mediated endocytosis, and regulation of the complement system (***Fig 6d***), whereas the cytosolic fraction had an upregulation of pathways associated with cell motility and response to unfolded protein (***Fig 6e***).

**Figure 6.**
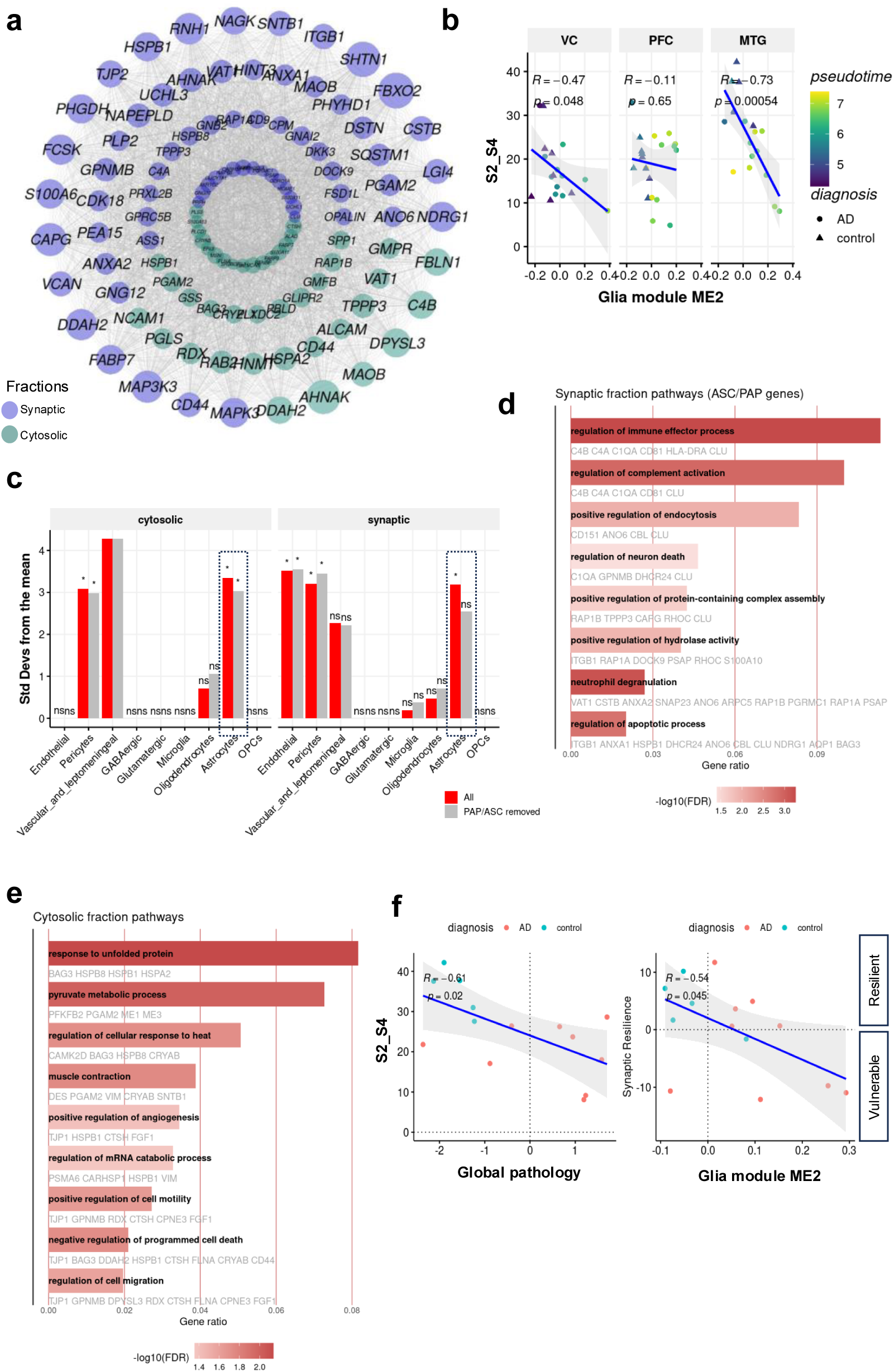
Astrocyte proteome enrichment at the synapse is dominated by astrocyte junction (ASC) and peri-synaptic astrocytic process (PAP) proteins. **(a)** Schematic co-expression network diagram showing module membership of protein from the cytosolic fraction (green) and protein from the synaptic fraction (purple) in the astrocyte module. Node size is proportional to connectivity (node with a degree >50 shown on the graph for clarity), and nodes are separated into four quantiles representing each layer. **(b)** Linear regression of summed density of S2 and S4 synapse subtypes (S2_S4) as a function of the glia module eigenvalue. **(c)** EWCE cell type enrichment of ME2 Glia module, stratified by cytosolic and synaptic fractions. Red bars indicate enrichment results on the complete cell-type data reference from the Allen Human Brain Atlas^59^. Grey bars indicate enrichment results on the same reference data but where PAP^61^ and ASC^60^ genes were filtered out. Enrichment p-values adjusted for multiple testing via Bonferroni correction (*ns>0.5, *p≤0.05, **p≤0.01, ***p≤0.001*). (**d,e)** Bar plots of the functional enrichment analysis against GO biological process terms on the synaptic fraction **(d)** and cytosolic fraction **(e)** proteins. Gene ratio represents the ratio of significant genes versus all annotated genes per pathway (colour: log_10_(FDR) one-sided over-representation Fisher’s exact test). **(f)** Astrocyte reactivity at the synapse worsens S2_S4 synaptic resilience. (Left) Scatter plot depicting the relationship between S2_S4 synapse density and global pathology (sum of log10(PHF1) and log10(amyloid)) in the MTG region, with a fitted regression line representing expected synapse density as pathology increases. This line serves as the baseline for synaptic resilience calculation, against which deviations are assessed to determine resilience scores. (Right) Scatter plot showing the relationship between synaptic resilience (observed minus expected S2-S4 synapse density) and UP_ME2 AD peri-synaptic astrocyte signature values.

NicheNet analysis was used to determine ligand-target matrix interactions between neurons and astrocytes converging at the synapse. The ligand with the potential to regulate the most targets was the upregulated DKK3, likely acting through its receptor KREMEN1 (***Supplementary Fig 10a-c***), both of which have been implicated in synapse loss^62,63^. NicheNet analysis provided a number of potential synaptic targets which DKK3 are likely to impact. These included proteins previously associated with synapse loss such as C3^64^, CCL2^65^, MMP9^66^, ELMO1^67^ and MMP7^68^.

Having related our AD peri-synaptic astrocyte signature to overall astrocyte reactivity, we wanted to study its relationship to synaptic resilience, that is, to assess whether the extent of synapse loss could be related to the extent of astrocyte reactivity at the synapse. S2 and S4 subtypes were previously defined as those with preferential loss in AD, specifically in the MTG region. We explored the relationship between S2_S4 summed density and global pathology (as the sum of log10(PHF1) and log10(Aβ) loads) in the MTG to derive a slope of expected synapse density according to pathology levels. A synaptic resilience score was then derived as the difference in observed minus expected synapse S2-S4 density (fitted values on the line) and related to UP_ME2 AD peri-synaptic astrocyte signature values. In this model, higher astrocyte reactivity at the synapse was associated with worse-than-expected synaptic resilience (or higher synaptic loss than predicted from global pathology). Altogether, GluN1+ synapse subtypes S2_S4 loss in the MTG could be worsened by astrocyte reactivity, suggesting a detrimental role at the synapse (R= −0.54, p-value= 0.045) ***(Fig 6f)***. Taken together, these provide a basis for understanding the intricate interactions and signalling pathways between peri-synaptic astrocytes and synapses in AD.

## Discussion

Synapse loss is the greatest neuropathological correlate to cognitive impairment in AD^69,70^ however, we are yet to determine how synaptic dysfunction, and subsequent loss progresses in the disease. Many studies, including our own^35,36^, have focused on synaptic changes in preclinical models in order to derive a longitudinal profile of synaptopathy but we lack an understanding of synapse vulnerabilities and of the molecular mechanisms that drive progressive synaptopathy in AD. Here, we have conducted a proteomic and synaptome study using multiple brain regions from AD and non-diseased control human *post-mortem* brain tissue to build a pseudotemporal profile of synaptopathy and identify vulnerable synaptic components. We identified potential therapeutic mechanisms by determining synapse-specific changes and elucidating the role of peri-synaptic astrocytic processes in synaptopathy.

### A pseudotemporal profile of synaptopathy in AD

The loss of synapses is a well-documented hallmark of AD, yet the events preceding synaptic loss remain relatively unexplored, especially in human AD. Our study aimed to characterize these early synaptic changes using synaptic proteomics and synaptome mapping across multiple brain regions in *post-mortem* AD tissue, representing region based sub-stageing of the disease^38,39^. The findings revealed that most differentially expressed proteins were brain region-dependent, with minimal overlap, indicating distinct regional differences. In the visual cortex, the region that experiences the shortest duration of disease, we observed evidence of metabolic downregulation, including mitochondrial dysfunction, a well-known early-stage AD characteristic^71^. Interestingly, we also detected upregulation of exocytosis, possibly due to increased presynaptic vesicle release due to mitochondrial dysfunction increasing the number of vesicles in the presynaptic active zone and reducing the threshold for vesicle release^72^. Early mitochondrial changes have also been implicated as precursors to hyperexcitability^73^, which may be an early consequence of synaptic dysfunction in AD. In the prefrontal cortex, a region typically affected at mid-stage AD^39^, there was a notable downregulation of inhibitory synapse assembly, evidenced by reduced levels of GABAergic receptor subunits. This finding aligns with the emergence of hyperexcitability in AD^74^. The PFC also exhibited downregulated mitochondrial function and upregulated exocytosis pathways, suggestive of continued elevated vesicle release, further contributing to hyperexcitability. In the region most affected by AD pathology, characterized by the greatest duration in the disease state, we found broad reductions in canonical synaptic transmission pathways and a decrease in GluN1+ synapse subtype density. This is consistent with previous findings of reduced PSD-95 levels in the temporal cortex and disruption of the excitatory-inhibitory balance and associated with cognitive impairment in AD^75^. Alongside distinct regional signatures, brain region-independent protein changes were also seen, with progressive differential expression across the visual, prefrontal, and temporal cortices. These included upregulated astrocytic markers and downregulated presynaptic scaffolding proteins. The correlation of those proteins with amyloid precursor protein levels and pathology (Aβ and phosphorylated tau accumulation) suggests they may serve as valuable biomarkers for tracking disease progression and evaluating therapeutic interventions. Collectively, these findings suggest a cascade of synaptic events across the disease spectrum, highlighting the progressive impact on the health of remaining synapses.

A key strength of this study is the direct comparison of multiple brain regions, which allowed us to hypothesise that our results represent a pseudotemporal profile of synaptic changes over the disease progression timeframe. We also derived a pseudotime measure based on AD-relevant factors and Braak staging, enabling a proxy analysis of synaptic changes during disease progression. However, the cortical regions assessed represent areas of distinct cortical functions and cellular organisation across the rostro-caudal axis, from high-level cognition in the PFC^76^, to complex visual and auditory object processing in the MTG^77^, and fundamental sensory input processing in the VC^78^. It is therefore possible that differences seen in the synapse proteome is partly a result of region-specific neuronal populations being vulnerable to pathology.

### Differential synaptic compartment and subtype changes in AD

Synapses are intricate structures composed of various components that develop and mature over time^79^. The use of untargeted proteomics in this study enabled a comprehensive assessment of synaptic changes in AD, revealing specific alterations that might be overlooked by more targeted approaches such as imaging. Given that synaptic function relies on both pre- and post-synaptic components, we examined the magnitude of differences between AD and control groups within each compartment. Our findings indicate that pre-synaptic proteins are affected earlier in the disease process, particularly in the VC region, while post-synaptic changes become more prominent as the disease progresses, especially in the PFC and MTG regions. This progression of synaptic changes in human AD is in contrast to preclinical models where post-synaptic components appear more resilient^35,36^.

While direct measurement of synaptic maturity in *post-mortem* tissue is not possible, unlike in preclinical models^35,36^, synaptic maturity can be inferred from synaptic NMDA receptor subunit protein composition. Through synaptome mapping, we identified distinct populations of excitatory GluN1+ subtypes of synaptic puncta. Our analysis revealed not just a loss of synapses but striking differential subtype vulnerability. We showed that specific subtypes of excitatory synapses were maintained while others were lost or gained in the MTG of late-stage AD cases. Specifically, synapse subtypes with the highest relative GluN1+ protein concentration (i.e. fluorescent intensity), indicative of more mature synaptic puncta on ‘mushroom’ dendritic spines^80^, are more likely to be lost in AD. This loss is consistent with preclinical findings that report reduced proportion of persistent dendritic spines^35^. Interestingly, we also observed an increase in puncta with immature properties (i.e. lowest relative GluN1+ protein concentration possibly on ‘thin’ dendritic spines) in AD, suggesting elevated synaptic turnover. Notably, we found an overall increase in the size of puncta, likely driven by the loss of the smaller higher intensity GluN1+ puncta, consistent with other studies^69^. This observation aligns with findings from preclinical tauopathy and amyloidopathy models^35,36^ and offers novel insight into the selective vulnerability and potential resilience or compensatory response of diverse synapse populations^81^. These results underscore the need for synaptotherapeutics to focus not only on synapses as a whole but also on specific compartments, subtypes, and the maturity of synapses, possibly down to the individual protein level. These findings also point to the need for further research to better understand the relationship between synaptic proteomic changes and their spatial distribution across different synaptic subtypes, which could have significant implications for the development of targeted treatments for AD.

The variability in the quality of human tissue presents challenges, particularly in the immunohistochemical staining of synaptic markers. In our study, due to inconsistent staining of synaptic markers across different tissue samples, only GluN1 proved to be a reliable marker for synaptome mapping ***(Supplementary Fig 13)***. This limitation restricted our assessment to the spatial distribution of GluN1+ puncta, representing one NMDA receptor subunit on glutamatergic excitatory synapses. GluN1 is considered the ‘obligate’ NMDA receptor subunit^82^ and so we considered it to be present in all excitatory synapses. However, there is a possibility that receptor subunit composition has changed. Other subunits showed no differential expression (GluN2a) or were also down-regulated (GluN2b) with GluN1 providing evidence for a loss of GluN1 synapses without a compensation of other NMDAR subunits. Consequently, the relationship between the proteomic features and spatial distribution of other synapse subtypes, such as inhibitory synapses, remains unclear. Given that proteomic changes in inhibitory synapses were more pronounced in the PFC, it is plausible that there would be a significant reduction in inhibitory synapse density or inhibitory interneurons in regions affected earlier in the disease, consistent with findings from other studies^26^. Additionally, as GluN1 is predominantly a post-synaptic marker, the density of pre-synaptic glutamatergic terminals may vary, particularly considering the differential pre- and post-synaptic proteomic changes observed in this study and in animal models^35,36^.

### The impact of peri-synaptic astrocytic processes in AD

The integration of data from synaptic and cytosolic fractions allowed us to further investigate the autonomy of synapses in the context of AD. We identified a synaptic module, predominantly driven by the synaptic fraction, which decreased as the disease progressed. This decline in the synaptic module aligns with worsening synaptic health and synapse subtype loss, rather than suggesting resilience in the remaining synapses. Additionally, the synaptic fraction was enriched with PAP^83^ proteins, such as Ezrin^84^, and led to the identification of a glial module—driven by both synaptic and cytosolic fractions—that increased with disease progression and was associated with synapse subtype loss^85^.

We identified an astrocytic signature at the synapse, which was associated with the loss of GluN1+ synapses, particularly in the middle temporal gyrus region. This signature included upregulation of endocytosis-related proteins, necessary for astrocytic neurotransmitter uptake^86^. Since PAPs are typically found at smaller synapses, astrocytic coverage—and thus neurotransmitter uptake—tends to be inversely correlated with synapse size^87^. We observed an increase in synapses with low GluN1 intensity, indicative of smaller synapses^88^ and possible greater coverage by PAPs which could lead to increased astrocytic uptake of glutamate. While this might help mitigate excitotoxicity, it could also impair efficient neurotransmission by reducing the available glutamate to spillover to neighbouring synapses^89,90^ or inhibit GABA release^91^. The astrocytic signature also included upregulation of complement activation components, such as C1Q and C4, which are implicated in astrocytic engulfment of synapses^92^. C1Q is involved in the initiation of complement cascades that lead to synapse pruning usually associated with microglia, while C4, primarily secreted by astrocytes^93^, is known to be increased in AD and associated with synaptic engulfment by astrocytes^94^. Furthermore, we saw an increase in MFGE8, with an emerging role as a linker between synapses and engulfing microglia^95,96^, but only in the PFC, the region with early signs of synapse loss. The greater astrocyte reactivity was associated with worse-than-expected synaptic resilience or greater synapse loss possibly by a greater synaptic engulfment or reduced stability of existing synapses. Additionally, the astrocytic module featured DKK3, a Wnt antagonist elevated in AD that contributes to the loss of excitatory synapses^97^, likely through interactions with KREMEN1—a pathway known to exacerbate synapse loss when dysregulated^62^. This analysis also provided a list of potential therapeutic targets by which the DKK3-KREMEN1 interaction might exert its effects on synapses. These findings provide evidence for a synaptotoxic astrocytic signature associated with PAPs in AD. However, elements of the astrocytic module, such as Clusterin^98^ and GPNMB^99^, have been associated with protective responses to synaptic damage, suggesting a partial protective response within the astrocytic environment. The neurotoxic astrocytic signature was primarily localized to the synaptic fraction, while the cytosolic proteins within this module were mostly linked to processes such as cell motility^100^ and responses to unfolded protein stress, consistent with the characteristics of reactive astrocytes that lose their supportive role for synapses in AD^101^. This highlights the role of the tripartite synapse as a distinct cellular compartment that is differentially affected in disease compared to the broader cellular environment.

As our *post-mortem* human tissue samples were obtained from later disease stages (Braak V-VI), it limits our ability to fully capture the true longitudinal course of disease progression, leaving open the question of whether the remaining synapses, particularly in the MTG where significant synapse loss was observed, are resilient or would eventually succumb to the same fate if disease progression had continued. Based on the synaptic and accompanying astrocytic signatures, we propose that the remaining synapses are in the process of being lost rather than exhibiting resilience. However, this hypothesis requires empirical testing to confirm. The SynPER synaptoneurosome preparation method used in this study enriches for, rather than purifies, synapses. This approach allows for the assessment of the tripartite synapse, including contributions from astrocytes, which is not possible with traditional synaptosome purifications. However, it also raises the possibility of contamination from non-synaptic components. To address this, we firstly characterized our synaptic proteome using a published synaptic protein exclusion list^12^, and secondly showed an enrichment for PAP/astrocytic proteins in our astrocytic module within the synaptic fraction when considering all proteins. While this does not completely rule out astrocytic contamination, it provides evidence that astrocytic proteins involved in the tripartite synapse are altered in AD and are associated with synaptic pathology.

Many omics studies in *post-mortem* brain tissue use single-nucleus RNA sequencing (RNA-seq) to understand differential responses in multiple cell types, but this approach has limited utility for studying synapses, as the cell body and nucleus are spatially distinct from neurites and synaptic compartments. Conversely, analysing synaptic and cytosolic fractions allows for the study of synapse-specific changes but presents challenges in assigning these changes to specific cell types. In this study, cell-type annotation against a reference database was used to attribute changes to neuronal or glial populations; however, broader cellular changes may be more difficult to assign to individual cell types. Moreover, fewer cytosolic proteins were identified (∼3000) compared to synaptic proteins (∼5000) and are more likely to be soluble rather than membrane-bound, however membrane-bound proteins were found in the cytosolic fraction (NCAM1, ALCAM).

The mechanisms behind the proteomic changes described here were not determined and it is imperative for these to be understood for synaptotherapeutic development. Most synaptic proteins are synthesised locally^102^ however this relies on the mRNA being available at the synapse. As nuclear transcription^103^ and cellular transport^104^ are affected by AD pathology, it is likely that mRNA is not available for local translation. Likewise, local protein degradation occurs at the synapse and is required for synaptic plasticity^105^ but this relies on a functioning ubiquitin-proteasome system. Proteasome function has been implicated in AD pathology^106^ which may extend to synaptic proteasome function. Taken together, these are two mechanisms by which the synaptic proteome may be affected in AD and warrant further investigation.

## Conclusion

In conclusion, this study provides evidence for impaired synaptic health in the remaining synapses and suggests a neurotoxic PAP phenotype in AD. These findings are suitable for hypothesis generation and target identification, and warrant further investigation in preclinical or cellular models to better understand the mechanistic underpinnings of synaptic and astrocytic interactions in the progression of AD. By building on previous research that characterised the synaptic proteome in AD, our findings provide novel insights into the time course and specificity of synaptic changes, and the role of astrocytes in synaptic pathology. These uncover new avenues for therapeutic interventions aimed at alleviating cognitive impairment in neurodegenerative diseases.

## Methods

### Subject selection

Samples were obtained with consent from three UK tissue brain banks; Edinburgh Brain Bank (EBBAD), London Dementia Brain Bank (LDBB), and South West Dementia Brain Bank (SWDBB). Fresh frozen brain tissue for synapse proteomics and formalin fixed paraffin-embedded (FFPE) tissue for synaptome mapping and pathohistological examination was sampled from the middle temporal gyrus (MTG, BA20/21), prefrontal cortex (PFC, BA9) and visual cortex (VC, BA17). A total of 13 Braak VI AD, and 13 sudden control cases were obtained across the three brain banks, a subset of which were used for synapse proteomics, synaptome mapping, and pathohistological examination (***Supplementary Data 1***). No major clinical co-morbidities (diabetes, symptomatic atherosclerotic disease or stroke, vascular disease, Lewy body or TDP43 pathology) were reported in case and control. Due to availability and high numbers of cases presenting with cerebral amyloid angiopathy (CAA) and hypertension, these were included in the selection. Subject demographics are outlined in ***Supplementary Data 1***.

### Histopathology

IHC was performed on 8μm FFPE sections from the MTG and PFC of each brain studied. Standard immunostaining procedures as recommended by the manufacturer were followed using the Super Sensitive Polymer-HRP (Biogenex) kit for β-Amyloid (4G8; Biolegend, 800711; 1:15,000) and pTau (PHF1; The Feinstein Institutes for Medical Research; 1:5000). Briefly, after dewaxing and rehydration of slides, endogenous peroxidase activity was blocked with 0.3% H_2_O_2_, followed by antigen retrieval (steamer, citrate buffer, pH 6.0). Primary antibodies were incubated overnight at 4 °C. Super Sensitive kit and DAB were used for antibody visualisation. Tissue was counter-stained by incubation in Mayer’s haematoxylin (TCS Biosciences) for 2 min, followed by dehydration, clearing and mounting. Digital images were generated from IHC stained slides scanned using a Leica Aperio AT2 Brightfield Scanner (Leica Biosystems). Images were analysed using Halo software (Indica Labs) after optimisation of Indica Labs macros for each antibody (β-Amyloid, PHF-1 = Area Quantification v1).

### Synaptome mapping: immunohistochemistry

FFPE 8μm sections were dewaxed and dehydrated through a series of xylene and graded ethanol solutions followed by an incubation in saturated picric acid to remove formalin sedimentation. Antigen retrieval was performed in citrate buffer (pH 6.0) in a decloaking chamber at 110°C for 15 minutes. Sections were then rinsed in ddH_2_O, washed in PBS, and blocked in solution containing 5% BSA, 0.5% Triton X-100 in 1xTBS for 1 hour at room temperature. Sections were then incubated overnight at 4°C for 18 hours with anti-GluN1/NR1 (IgG1, Mab, 75-272, 1:250 (0.004mg/ml), Neuromab) in antibody solution containing 3% BSA, 0.2% Triton-X in 1X TBS. The next day, primary antibody solution was washed off using 1X TBS with Triton X-100 before a 2-hour incubation with Cy5-AffiniPure Goat Anti-Mouse IgG, Fcγ Subclass 1 (1:250 (0.0048mg/ml), 115-175-205, Jackson ImmunoResearch) in antibody solution. Finally, sections were counterstained with 4′,6-diamidino-2-phenylindole (1 μg/ml DAPI (Sigma) diluted 1:1,000 in PBS) and mounted on to coverslips using mounting medium (mowiol with 1,4-diazabicyclo[2.2.2]octane (DABCO), Sigma-Aldrich).

### Synaptome mapping: image acquisition, extraction, and delineation

Single-synapse resolution imaging of GluN1+CY5+ synaptic puncta was achieved using an Eclipse Ti2-E Nikon spinning-disk confocal microscope equipped with a CSU-W1 (pinhole size 50µm), a CFI HP Plan Apochromat VC 100X oil-immersion lens (NA 1.40), and a Photometrics PRIME BSI camera. Image tiles of 942 x 920 pixels (64.42 x 62.92 µm) in size and 16-bit depth were acquired. To cover large regions of interest, control X, Y, and Z points were inputted around the circumference of the tissue, generating a focus surface for the acquisition of multi-tile single plane nonoverlapping images. To correct for focus drift over a large area, the focus surface was recalculated every 400 tiles and the focus surface position adjusted accordingly. Individual tiles from raw nd2 image files were extracted as TIFF files along with image metadata. Image tiles were then stitched to generate overview montage images that were downsized by a factor of 16 using a custom MATLAB script. Cortical layers and white matter were manually delineated on Nissl-stained serial sections using the polygon selection tool in FIJI which were subsequently overlaid on to the overview montage images and finalised before being saving as a set of .roi files.

### Synaptome mapping: synaptic puncta detection and analysis

GluN1+CY5+ synaptic puncta were detected and segmented from the acquired fluorescent tile images by developing a Mixture of Experts ResUNet model. The model was trained using a manually annotated training dataset taken from 5 sudden death control and 5 late-stage AD human *post-mortem* cases. Sub-tiles (128 x 128 pixels) from a training set of 1674 images were randomly sampled across layers and subregions from 3 x cortical (frontal, temporal, entorhinal), 1 x subcortical (putamen), and 1x hippocampal brain regions. All puncta in the training set were manually annotated by 5 independent scientific investigators. Annotation of each punctum was weighted per individual and the ground truth was generated by weighted averaging over all individuals with a true punctum being determined with an average weight greater than 0.7. Ensemble learning was applied for training and a K-folder method was used to validate and test the trained classifier.

After puncta were detected and segmented, puncta parameters (mean pixel intensity, size, skewness, kurtosis, circularity and aspect ratio) were calculated based on the estimated full width half maximum of each punctum and the puncta density per unit area was quantified. Based on the puncta parameters measured, GluN1+CY5+ synaptic puncta were classified into 4 subtypes using an unsupervised Mixture of Experts machine learning model. These 4 subtypes were mapped across AD and control single-synapse resolution images and the average subtype density was quantified and averaged across all cortical layers in a 6mm x 6mm window, excluding delineated white matter regions.

The Shapiro-Wilk test was performed to evaluate the normality of the data distributions, and Levene’s test was used to assess the homogeneity of variances. Based on these results, unpaired, two-sided t-tests were conducted for AD and control comparisons where both the normality and homogeneity of variance assumptions were met. For comparisons where either assumption was violated, the non-parametric two-sided Mann-Whitney U test was used. For the GluN1 subtype analysis, due to multiple subtype comparisons, p-values were adjusted using the Benjamini-Hochberg procedure to control the false discovery rate. *Post-mortem* interval (PMI) did not affect puncta density (***Supplementary Fig 13c***).

### Synaptic and cytosolic protein extraction

Synapse-enriched and cytosolic fractions were isolated from homogenized human cortical tissue using the SynPER (Synaptic Protein Extraction Reagent) preparation protocol as per the manufacturer’s instructions. Approximately 100mg frozen tissue samples were thawed and homogenised using 15 strokes of a dounce homogeniser in Synaptic Protein Extraction Reagent (SynPER, Thermofisher) and Complete Protease Inhibitor (EDTA Free, Roche). Following homogenisation, samples were centrifuged at 1,200g for 10 minutes at 4°C. The pellet was discarded, and supernatant was transferred to a separate sample tube for centrifugation at 15,000g for 20 minutes at 4°C. The supernatant (cytosolic fraction) was extracted, snap frozen, and stored at 80 C. The remaining pellet (synapse-enriched fraction) was resuspended in cold PBS before a final centrifugation step at 15,000 x g for 20 minutes at 4°C. The supernatant was discarded, and the remaining pellet (synapse-enriched fraction) was snap frozen at stored at −80 C.

### Label-free quantitative proteomics

Synaptic proteins were eluted from synaptoneurosomes by the addition of 100μL 8M urea, with 10 minutes in a sonicating water bath at room temperature, followed by centrifugation for 10 minutes at 4°C at 10,000xg. Cytosolic samples were prepared in SynPER buffer. A protein assay was performed on all synaptic and cytosolic fractions according to the manufacturer’s instructions (DC Protein Assay microplate protocol, Bio-Rad). 20μg of protein were then taken from each sample and digested using an automated method on an Andrews+ pipetting robot (Andrews Alliance).

Briefly, samples were transferred to a digestion plate and held at 4°C while being combined with ddH_2_O. A digestion buffer was then added such that the final reaction conditions were 10mM tris(2-carboxyethyl)phosphine, 40mM chloroacetamide, 100mM ammonium bicarbonate, and 0.2μg trypsin in 120μL, ensuring the urea concentration was <2M across all synaptic samples. Samples were then heated to 37°C for 16 hours before the reaction was halted by reducing the temperature to 4°C and adding 0.5% (V/V) trifluoroacetic acid (TFA).

Samples were desalted using a 96-well Oasis μElution HLB plate (Waters), where the plate was first equilibrated with 3×100μL 0.1% TFA drawn through under negative pressure. Samples were loaded onto the plate and washed with 3×100μL 0.1% (V/V) TFA before being eluted with 150μL 60% (V/V) acetonitrile and dried completely in a SpeedVac (Eppendorf) at 60°C. Peptides were dissolved in 50μL 0.1% (V/V) formic acid (FA), and a peptide assay was performed as previously described. Concentrations of peptides were then adjusted to 0.06μg/μL for analysis by LC-MS/MS.

Peptides were analyzed by data-independent acquisition LC-MS/MS on an M-Class UPLC (Waters) coupled to a ZenoTOF 7600 (Sciex) in ZenoSWATH mode. 300ng were loaded onto a Kinetex 2.6μm XB-C18 150×0.3mm column (Phenomenex) for 1 minute and eluted over a 20-minute gradient, with the organic mobile phase increasing from 3% (acetonitrile with 0.1% (V/V) FA) to 30% at a flow rate of 10μL/min. After a column clean at 80% organic phase at 10μL/min for 2 minutes, the buffers returned to starting conditions of 97% aqueous phase (0.1% (V/V) FA) for 6 minutes. The column was heated to 50°C throughout.

The MS was run in positive mode, acquiring MS data between 400 and 1500Da with a 0.1s accumulation time. MS/MS data were acquired between 140 and 1800Da with dynamic collision energy enabled and an accumulation time of 0.013s. 85 SWATH windows were variably distributed between 399.5 and 903.5Da. The LC and MS were operated using OS software version 3.1 (Sciex). A study reference of pooled synaptic or cytosolic samples was run every eight samples, and a blank was run between each injection.

### Processing of proteomics data

Proteomics raw data was searched against a predicted library of human canonical proteins using DIANN (V1.8) and annotated with sequence information from Swissprot (Uniprot) downloaded 03/05/03. Match-between-runs was enabled. Bulk proteomics synaptic and cytosolic fractions were processed using the Omix R package V1.0^107^ to generate a comprehensive multi-assay dataset. The raw synaptic fraction comprised 6139 proteins, while the cytosolic fraction 3062. Briefly, proteomics processing was achieved via the *process_protein()* function from the Omix R package (v1.0). Proteins with missing values in more than 50% of samples were filtered out. After filtering, 5755 and 2528 proteins were kept in the synaptic and cytosolic fractions, respectively (***Supplementary Data 2-3***). Proteins were imputed following the minimum value method (*imputation=’minimum_value’*). A batch correction was applied using median centring (*correction_method=“Median_centering“*).

### Differential protein abundance

Differential protein abundance between AD and control groups was quantitatively assessed using the limma package (v.3.50.3)^108^. We used the *lmFit()* function with its robust regression option to construct linear models, accommodating biological variability and potential outliers.

In the regional AD versus control differential expression analysis, the model included age, sex, and PMI as covariates. Contrasts were defined and applied using the *contrasts.fit()* function, and empirical Bayes moderation was performed with robust variance estimation (*eBayes(vfit, robust = TRUE)*). P-values from the moderated t-tests were adjusted for multiple testing using the Benjamini-Hochberg procedure to control the false discovery rate (FDR)^109^. We applied a significance threshold of pval<0.05 and Log2FC threshold >0.2 for downsteam analysis. Sensitivity analyses included continuous regressions against neuropathological measurements of amyloid and pTau, to highlight similarities with case/control setup (***Supplementary Data 4***). This was performed only on the MTG and PFC regions as the VC regions lacked neuropathological measurements.

In the pooled AD versus control differential expression analysis, samples from all brain regions were pooled. The model included age, sex, PMI, and brain region as covariates. Differential expression analyses were performed separately for synaptic and cytosolic fractions. For sensitivity analysis, we used a linear mixed model with a random intercept (∼ diagnosis + age + sex + PMI + Region + (1|case_id)) to account for within-subject correlations. While proteins in the mixed model did not survive multiple testing, a 94% match was observed when comparing proteins with significant p-values from the mixed model to those from the pooled analysis. Overlapping proteins were further filtered for those differentially expressed in both fractions and compared for concordance in directionality.

### Functional enrichment

Functional enrichment analysis was performed using Enrichr^110^. This streamlined pathway analysis allowed for the generation of pathway reports utilising common pathway databases such as KEGG, Reactome, BioCarta, MSigDB, and Gene Ontology.

### SynGO enrichment

Regional differentially expressed proteins were analysed on the SynGo portal, with brain expressed proteins as background. SynGo enrichment plots were customised via http://syngoportal.org/plotter#. The Gene Ratio was derived as the ratio of GSEA count over GSEA background.

### Synaptic and cytosolic fractions integration

Integration of the synaptic and cytosolic fractions was performed using the Multi-Omics Factor Analysis (MOFA) model from the MOFA2 R package (v1.4)^57^. Before integration, scaling of the different fractions was carried out to ensure comparability. Datasets used for integration comprised the synaptic proteomics (4925 proteins), the cytosolic proteomics (2204 proteins), of which 2075 features overlapped between the two fractions fractions. To ensure precise protein identification, ambiguous entries representing multiple isoforms or homologous proteins (e.g., “C4A;C4B”) were removed from the dataset. Unique protein identifications, such as C4B and C4A, were retained for integration. The MOFA model was configured to extract ten latent factors, comprehensively representing the underlying biological processes. Default settings were used for other parameters.

Factor enrichment was performed using the *run_enrichment()* function with default parameters and SynGo ontologies^46^ as feature.sets. Factor 3 was chosen for downstream analysis, given it was enriched for synaptic gene ontologies and significantly correlated to AD neuropathology measures.

### Pseudotime derivation

To compensate for the lack of longitudinal tissue data in *post-mortem* samples, we employed a pseudotime trajectory analysis to model synaptic-cytosolic dynamics across a disease continuum. It uses the same concept as in pseudotime inference with scRNA-seq^111^, except here, each data point is a bulk sample derived from a patient, and pseudotime represents an inferred trajectory along which patient samples are ordered, modelling disease progression from early to advanced stages. The disease trajectory was derived using the slingshot algorithm^112^ on a two-dimensional UMAP representation of AD-relevant and brain region-specific MOFA factors (Factors 1,2,3). We clustered the latent space (UMAP1/UMAP2) into 3 groups via k-means clustering and chose the root of the slingshot trajectory as the cluster with a high density of VC samples (cluster 3) and ending cluster enriched in MTG samples (cluster 2) (***Supplementary Fig 8)***. This was specified in the slingshot parameters (start.clus =3,end.clus=2). We then ranked samples according to their position along the slingshot trajectory but within predefined Braak tiers (early [Braak 0-II] and late [Braak V-VI]), implementing a semi-supervised approach. The final pseudotime values were determined by fitting a linear model between the derived disease trajectory and the sample ranks within each Braak tier. This method allows us to model AD progression as a continuous process, positioning each patient along an inferred gradient of disease severity.

### Synaptic-cytosolic proteomics co-expression modules

Factor 3 was selected for co-expression module analysis. Factor 3 features with weights exceeding ±1.5 SD from the mean were used to define the signature. Since this factor is negatively correlated with clinical covariates, features with negative weights are expected to increase with disease progression (“*UP*”-labelled features). Conversely, features with positive weights are expected to down-regulate with disease progression (“*DOWN*”-labelled features). Pairwise correlations between features in the UP/DOWN signatures were assessed, and a synaptic-cytosolic co-expression network was constructed where feature pairs showed an absolute Pearson’s correlation exceeding a user-defined threshold (default=0.3). This network was then dissected into functionally relevant modules using community detection algorithms like Louvain^113^, resulting in disease-correlated modules that aid in understanding disease mechanisms. Module eigenvalues were calculated via Principal Component Analysis on scaled expression data, and the first principal component was assigned as the module eigenvalue.

### Targeted exclusion cell type enrichment

Cell-type enrichment analyses were conducted using Expression-Weighted Cell Type Enrichment (EWCE) (v1.2)^58^. This analysis utilised a cell-type data reference from the Allen Human Brain Atlas^59^ to determine the enrichment of specific cell types within the identified modules. To refine our understanding of astrocyte involvement at the synaptic level, we repeated the cell type enrichment analysis after excluding genes known to be involved in peripheral astrocyte processes (PAP)^61^ and neuron-astrocyte junctions (ASC)^60^.

### Gene set enrichment

To assess the statistical overlap between our results and ASC/PAP gene sets, we used Fisher’s exact test via the Gene Overlap package (v4.4). This test calculates a p-value to evaluate whether the observed overlap is likely if the gene sets are independent. It also provides an odds ratio, a measure of the association strength between the gene sets. An odds ratio of 1 or less indicate no association. We considered all expressed genes as the background set, following default settings.

### ASC/PAP activity

To determine ASC/PAP gene set activity in our synaptic proteomics data, we used the GSVA package (v1.42). GSVA is an unsupervised method that quantifies changes in pathway activity on a per-sample basis. We applied this analysis using the default parameters.

### Cell-cell communication analysis

Cell-cell communication analysis was performed using NicheNet (v1.1.1)^114^. We aimed to identify and prioritise ligand proteins in astrocytes at the synapse that could potentially influence synaptic proteomics changes. NicheNet uses gene expression information from the dataset under study and integrates this information with a prior model built by integrating prior knowledge on ligand-to-target signalling paths. NicheNet constructs a model from existing knowledge of ligand-target relationships, aiming to identify ligands produced by “sender” cells that could influence a specific set of genes in “receiver” cells. This influence extends beyond the immediate receptors to any relevant downstream gene targets. For this analysis, we used NicheNet’s default settings. The “receiver” cell gene set was defined to include proteins from the GluN1+ module and its interactome (see below methods). The “sender” genes and ASC and PAP proteins comprised the combined synaptic fractions of the ME2 glia/astrocyte module.

### Defining the GluN1+ interactome

We defined the GluN1+ (GRIN1) interactome using the STRING database (version 12.0)^115^, explicitly sourcing data from the “9606.protein.links.detailed.v12.0.txt” file. We focused on hub proteins in the synaptic fraction of the GluN1+ module and included only interactions with a minimum experimental score of 700, indicating solid experimental evidence. Interactions with mitochondrial proteins marked by the “MT-” prefix were excluded to prevent confounding effects.

### Synaptic proteomics genetic enrichment

We used MAGMA software (version 1.10)^116^ for genetic enrichment analysis of upregulated proteins. We extracted SNP locations and p-values from harmonised GWAS summary statistics^50–52^. Using these SNP locations and an NCBI gene location file (NCBI38.gene.loc), we annotated SNPs to genes using MAGMA’s --annotate command. We then analysed gene-based association with reference genotypes from the 1000 Genomes Project (European population). We calculated gene-based p-values using our gene annotation file and the GWAS summary statistics. This involved calculating gene-based association results separately for three datasets (Bellenguez_magma.genes.out, Janssen_magma.genes.out, and Wightman_magma.genes.out) and then combining these results with the --meta command to generate meta-analyses p-values (meta_analysis.genes.out). The enrichment analysis incorporated these individual and combined statistics, assessing the genetic associations of different upregulated genes using MAGMA’s --set-annot command.

## Data availability

The mass spectrometry proteomics data have been deposited to the ProteomeXchange Consortium via the PRIDE^117^ partner repository with the dataset identifier PXD056052.

## Acknowledgements

We thank the donors and their families for the use of human brain tissue in this study and the UK brain bank staff for making it available. Tissue samples were provided by the London Neurodegenerative Diseases Brain Bank at King’s College London. The brain bank receives funding from the UK Medical Research Council and as part of the Brains for Dementia Research programme, jointly funded by Alzheimer’s Research UK and the Alzheimer’s Society. We would like to thank the South West Dementia Brain Bank (SWDBB) for providing brain tissue for this study. The SWDBB is part of the Brains for Dementia Research programme, jointly funded by Alzheimer’s Research UK and Alzheimer’s Society and is supported by BRACE (Bristol Research into Alzheimer’s and Care of the Elderly) and the Medical Research Council. We also thank the Edinburgh Brain Bank for the provision of tissue and expert advice from neuropathologist, Professor Colin Smith. The provision of case data used in this study was provided with support from the BDR programme, jointly funded by Alzheimer’s Society UK and Alzheimer’s Society. We are grateful to Diana P. Benitez for her support in the human tissue management. We would also like to thank Digin Dominic for the generation of the synapse visualisation tool. The authors thank Aina Badia Soteras for critical evaluation of the manuscript. Supplementary figure 1 was created with BioRender.com.

Work in the SG laboratory was funded by the UK Dementia Research Institute which receives its funding from UK DRI Ltd (award XCPD2019–01) funded by the UK Medical Research Council, Alzheimer’s Society and Alzheimer’s Research UK (JG) and by Wellcome Trust grant 218293/Z/19/Z (BN, RQ).

PMM acknowledges generous personal support from the Edmond J Safra Foundation and Lily Safra and an NIHR Senior Investigator Award. JSJ was supported by funding from Alzheimer’s Society (grant number 599 (AS-DRL-22-008)) and Rosetrees Trust (PGS21/10048). This work was supported by the UK Dementia Research Institute, which receives funding from UK DRI Ltd., funded by the UK Medical Research Council, Alzheimer’s Society, and Alzheimer’s Research UK. The study is an output from the UK Dementia Institute Multi-omics Atlas Project for Alzheimer’s Disease (MAP-AD; map-ad.org). Infrastructure support for this work was provided by the NIHR Imperial Biomedical Research Centre and Medical Research Council World Class Labs Award (grant number MC_PC_MR/X013537/1)

For the purpose of open access, the author has applied a CC-BY public copyright licence to any Author Accepted Manuscript version arising from this submission.

## Author Contributions

All authors contributed, reviewed and approved the manuscript.

