## Supplementary Figures for "A synaptic-astrocytic proteomic signature associated with synaptopathy in Alzheimer’s Disease"

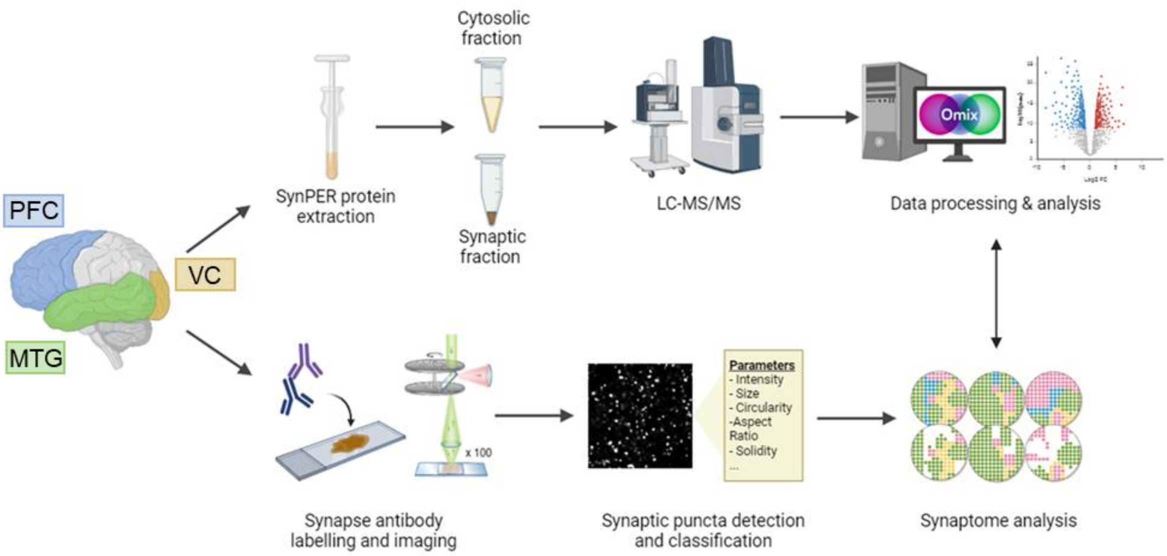


Supplementary figure 1: Synaptomics methodology. Schematic diagram of the experimental workflow. Samples were collected from the visual cortex (VC), prefrontal cortex (PFC), middle temporal gyrus (MTG), and middle temporal gyrus (MTG) from control and AD brains. Synapse enriched and cytosolic fractions were isolated using the SynPER synaptic protein extraction method. Both fractions underwent proteomic analysis using liquid chromatography coupled with mass spectrometry-based quantification and identification (LC–MS/MS). Proteomics data were analysed using Omix, a multi-omics' integration pipeline[1]. In parallel, formalin fixed paraffin-embedded (FFPE) sections from the same cases* were immunolabelled with the synaptic marker, GluN1, and imaged at single-synapse resolution. Synaptic puncta were detected, segmented, and classified into subtypes using synaptome mapping analysis. *Subset of cases used dependent on tissue availability.

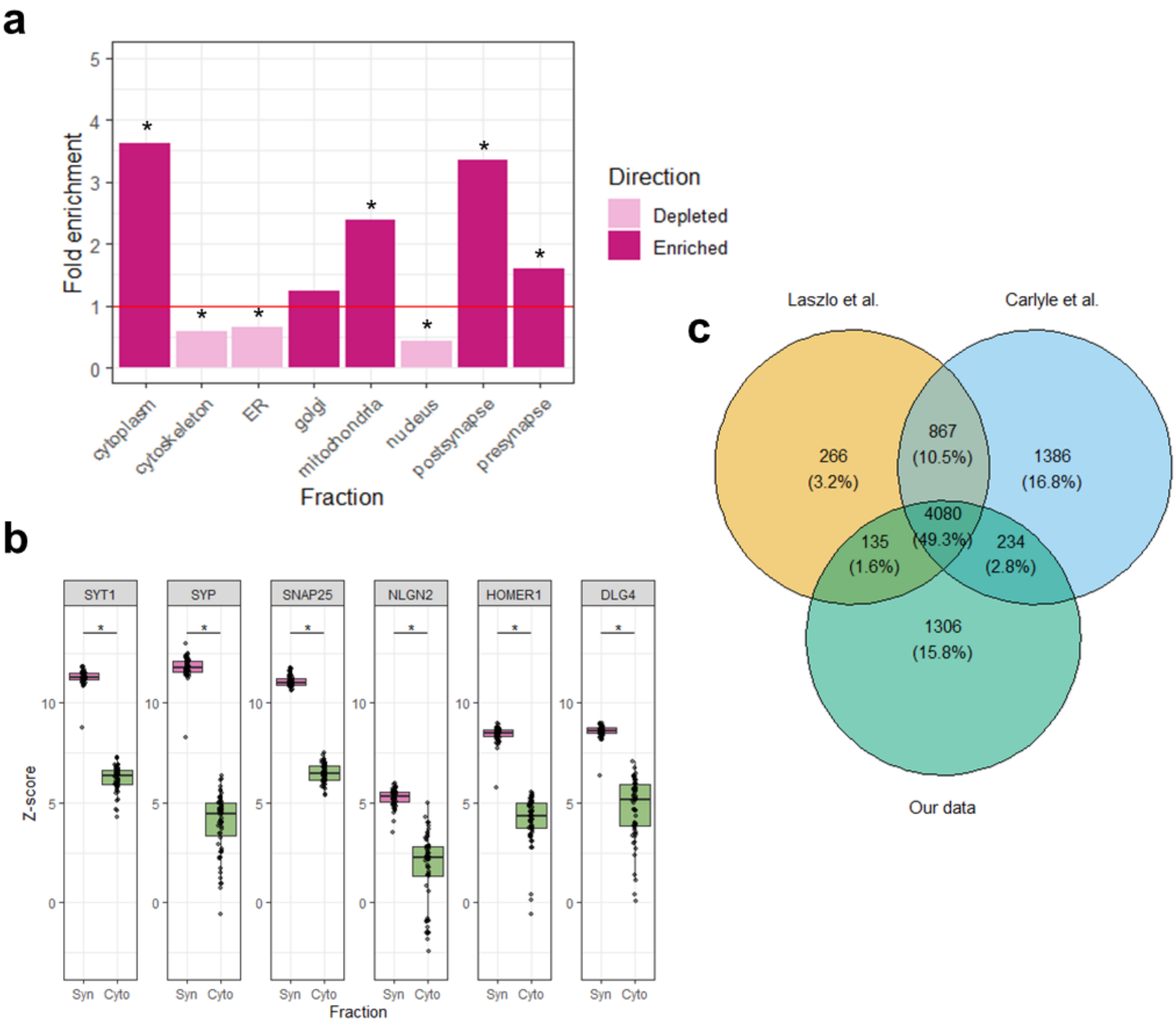


Supplementary figure 2: Synapse enrichment in human post-mortem cortical tissue. **(a)** Comparison of detected proteins in the synaptic fraction against a consensus list of unique cellular fraction-associated protein IDs (*Fisher's test, Bonferroni-adjusted p-value < 0.001). **(b)** Boxplots of the normalised abundance of selected pre- (SYT1, SYP, SNAP25) and post-synaptic (NLGN2, HOMER1, DLG4) proteins in synaptic fraction (Syn) compared to cytosolic (Cyto) (*p≤0.05, Mann-Whitney U test with continuity correction). **(c)** Venn diagram illustrating the intersection of 5755 uniquely identified proteins with previous human synapse proteomic datasets.

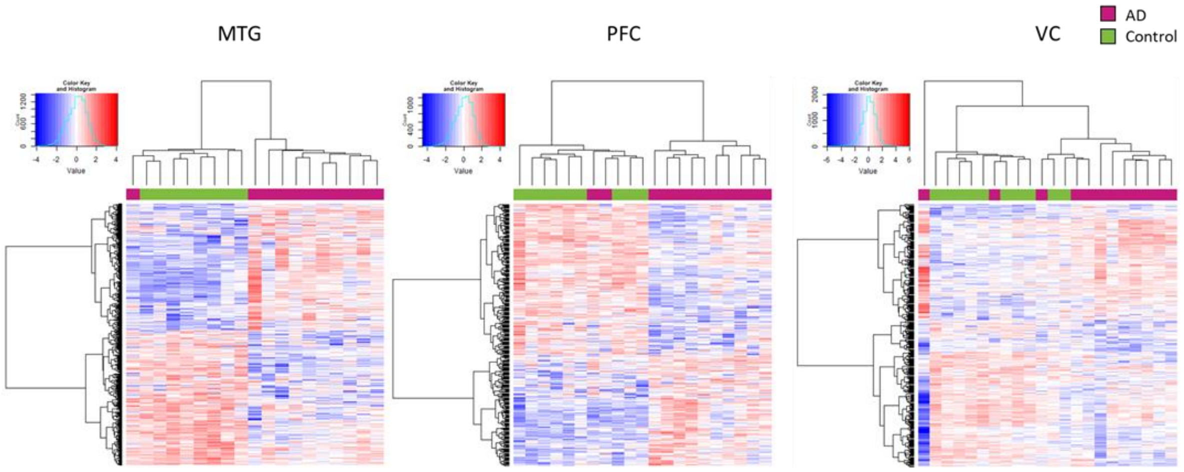


Supplementary figure 3: Relative abundance of differentially expressed proteins in the MTG, PFC, and VC distinguishes AD and non-disease control groups. Heatmaps of DEP abundance in AD and control cases in the MTG, PFC, and VC. Euclidean clustering based on abundance of DEPs and cases (colour: Z-score).

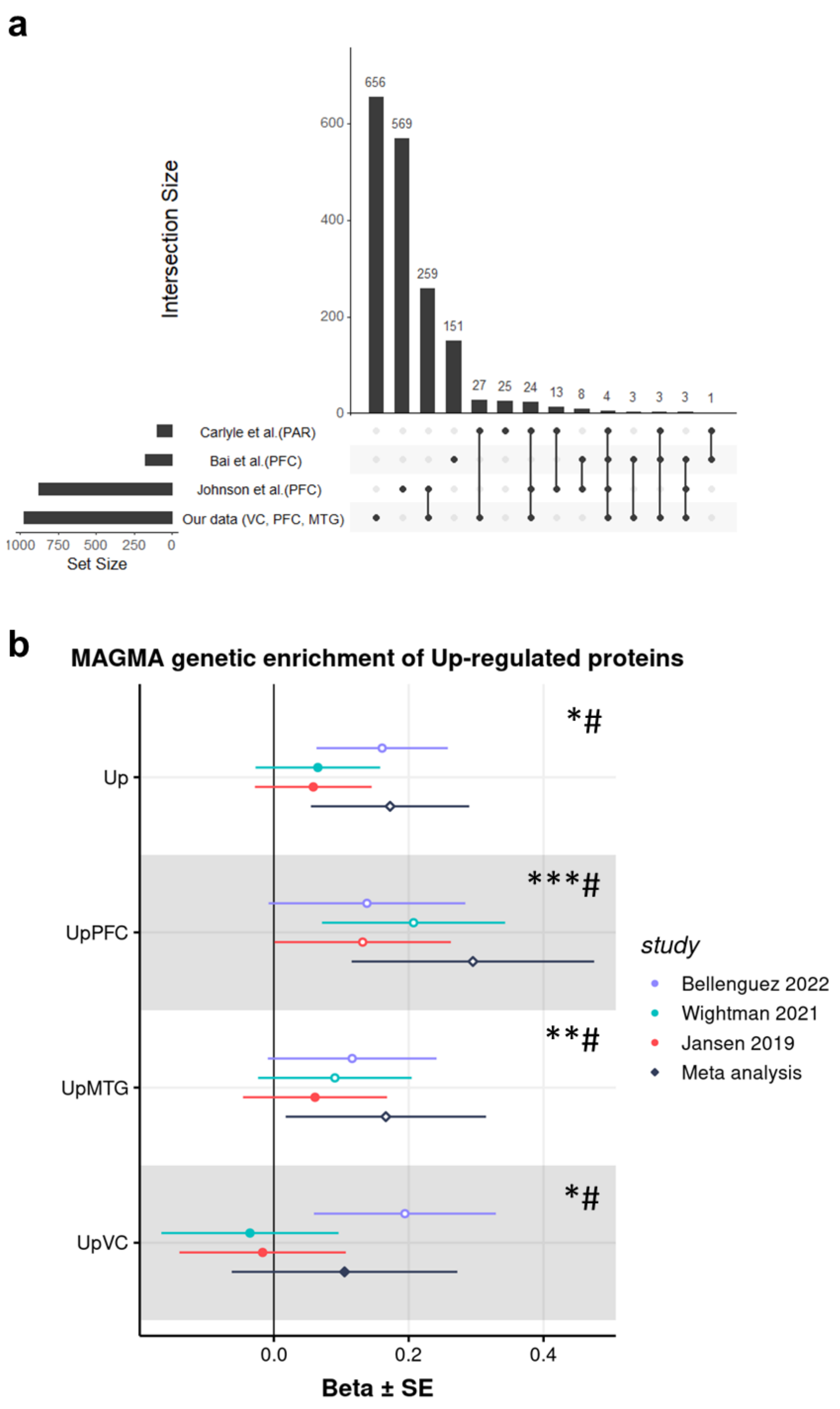


Supplementary figure 4: Study comparisons. **(a)** Upset plot showing the intersection of DEPs reported in other synapse[2] and bulk proteomic datasets[3,4], as well as our own. Brain region/s assessed in each study are marked in brackets next to study reference (PAR, parietal association cortex; PFC, prefrontal cortex; VC, visual cortex; MTG, middle temporal cortex). Full comparisons of each reported protein are detailed in ***Supplementary Data 6***. **(b)** Forest plot illustrating the enrichment of upregulated proteins (all regions, PFC, MTG, VC) in genetic risk quantified by three AD GWAS 6–8 and in a meta-analysis, highlighting significant beta values and corresponding P-values. # indicates modules enriched in AD risk genes in the meta-analysis and * only in one or more studies.

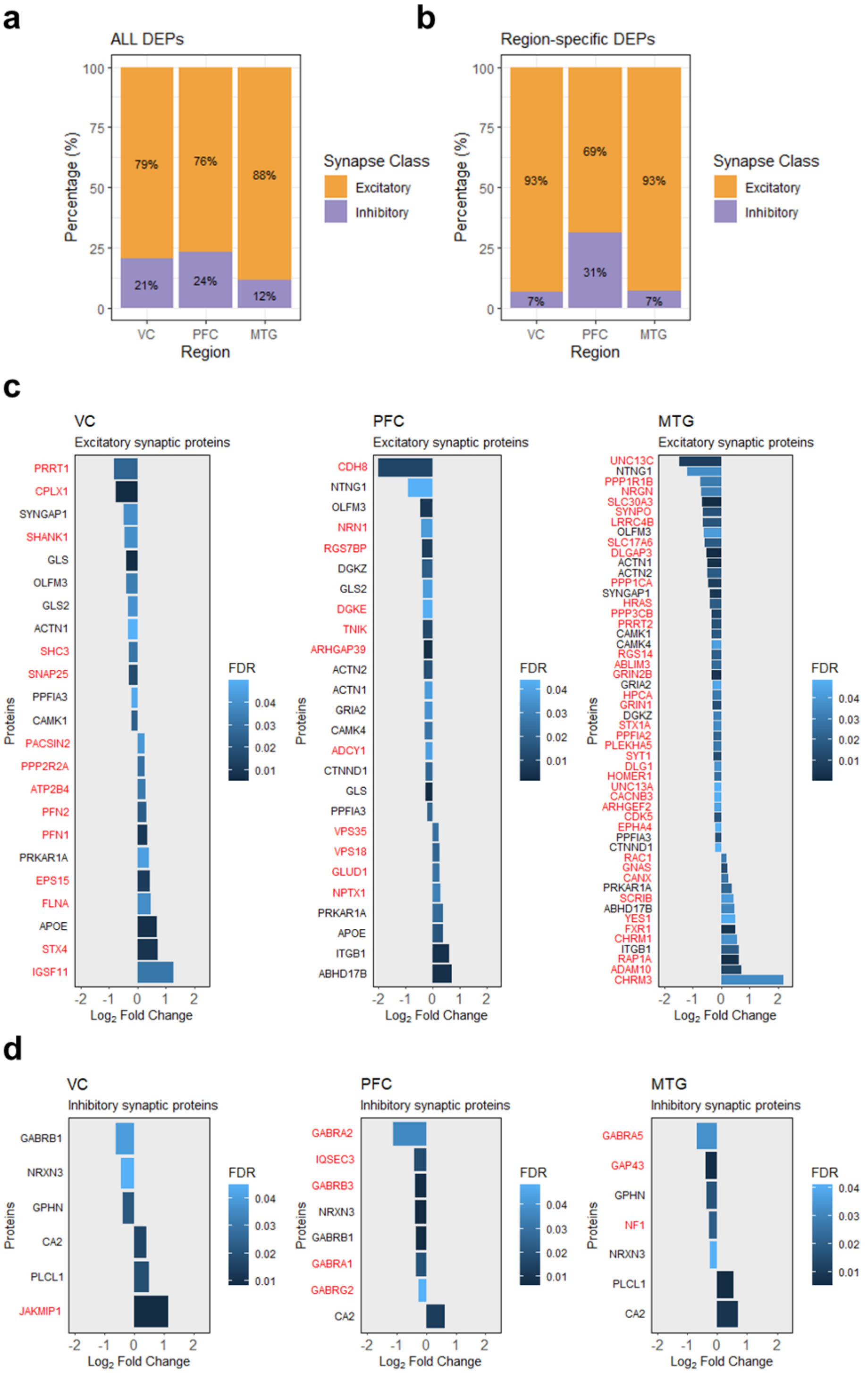


Supplementary figure 5: Regional excitatory and inhibitory synapse protein signatures differentially expressed with AD. **(a, b)** Stacked histogram displaying the percentage of DEPs found in excitatory or inhibitory synapse classes based on a consensus list of proteins[5]. **(c,d)** Barplots showing the $\log_2$ fold change of those excitatory **(c)** and inhibitory **(d)** DEPs per region, ranked in order of fold change. Region-specific DEPs are labelled in red (colour: FDR, one-sided over-representation Fisher's exact test).

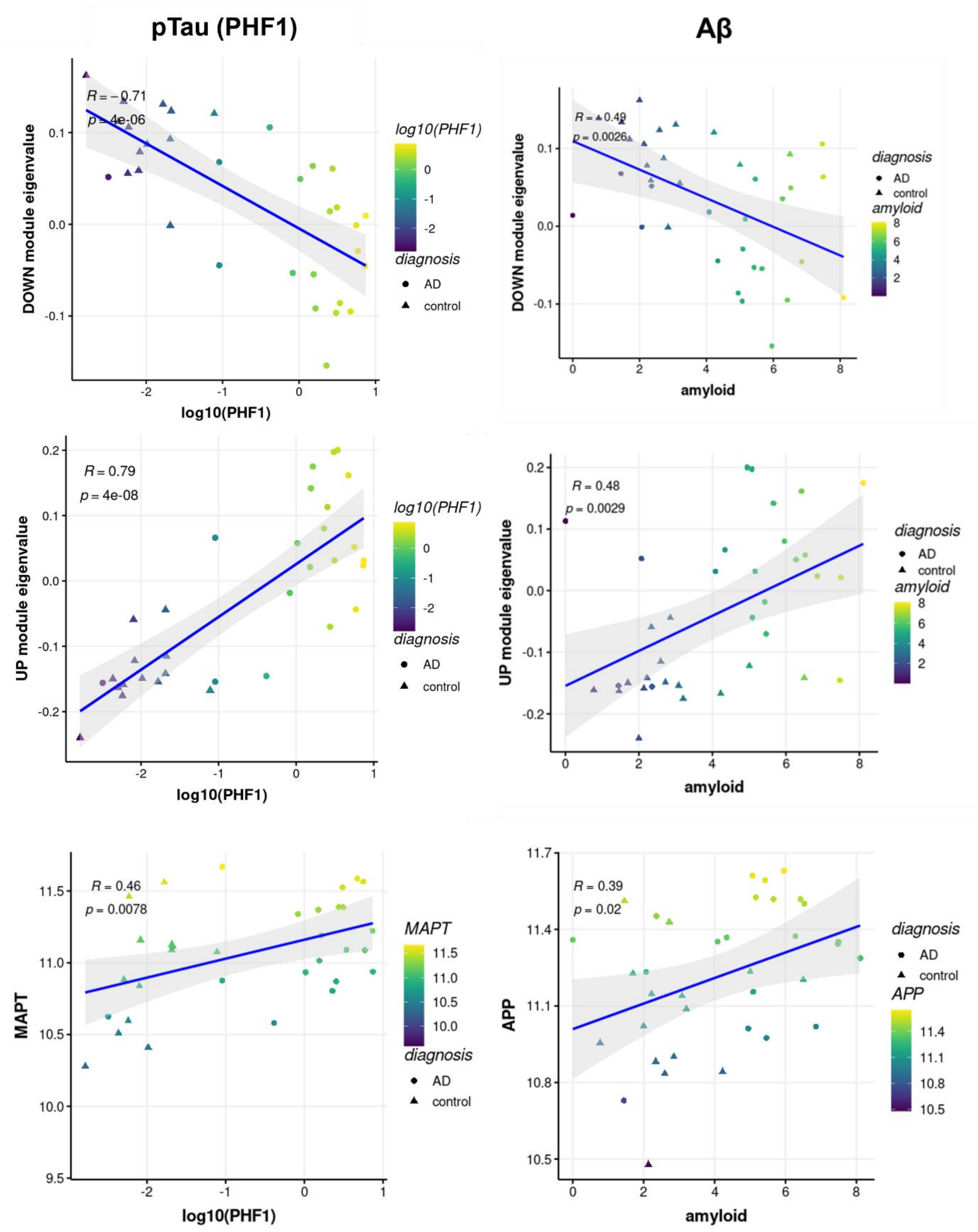


Supplementary figure 6: Brain region-independent DEPs correlate with AD pathology. Linear regression of the upregulated and downregulated module eigenvalues against pTau (PHF1) and Aβ (4G8, amyloid) percent immunoreactive area. Final row shows linear regressions against MAPT and APP protein abundances against pTau (PHF1) and Aβ (4G8, amyloid) percent immunoreactive area.

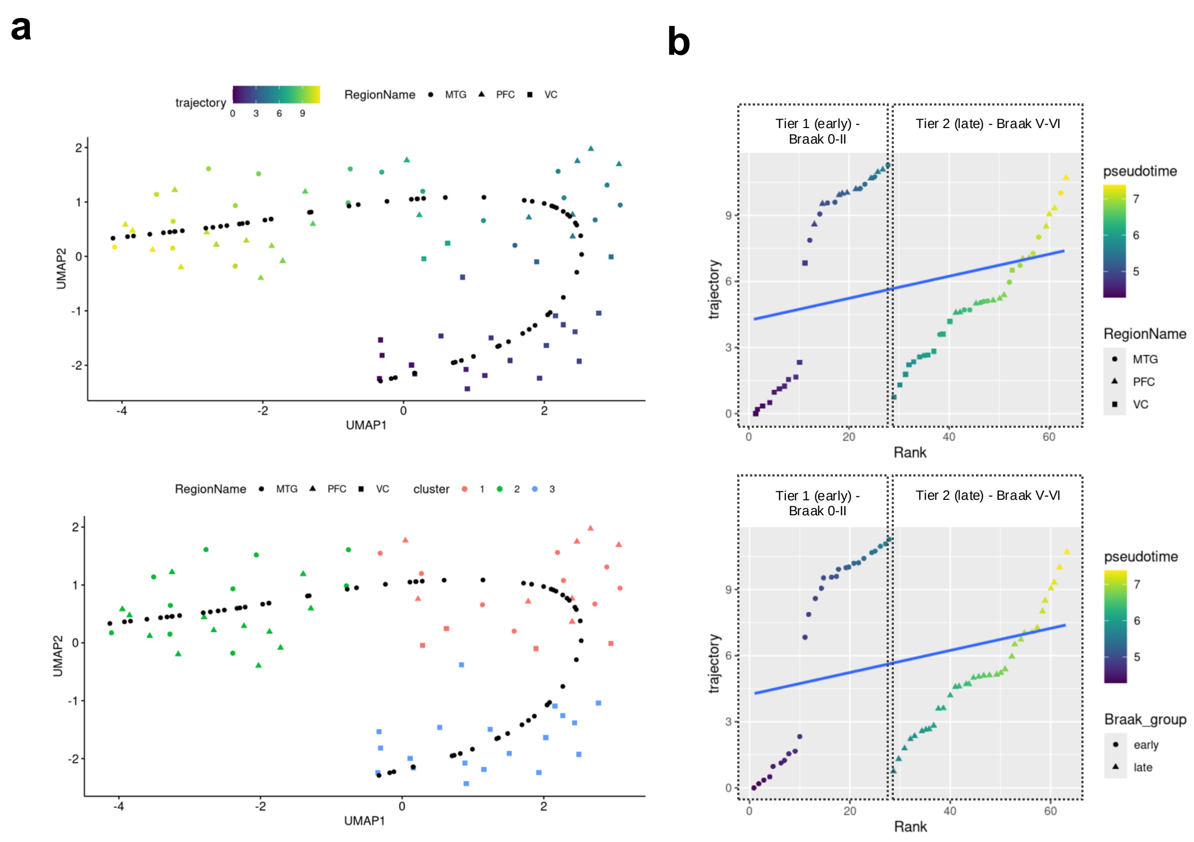


Supplementary figure 7: Semi-supervised approach for pseudotime derivation.

**(a)** AD-relevant factors 1,2, and 3 were used in a semi-supervised approach to derive a pseudotime measurement. This figure presents a Uniform Manifold Approximation and Projection (UMAP) plot illustrating the distribution of brain tissue samples based on inferred pseudotime (i.e. trajectory) derived via Slingshot[6]. Each point represents a tissue sample coloured by inferred pseudotime, ranging from early (blue) to late (yellow) stages of the disease. The VC dense region (cluster 3) was chosen as the root of the slinghshot trajectory. **(b)** This plot illustrates the inferred trajectory of AD progression plotted against the ranked positioning of brain tissue samples according to Braak staging criteria into tiers. The x-axis represents the overall rank of samples from early to late Braak stages, while the y-axis denotes the trajectory inferred through pseudotime analysis. The linear model depicts the relationship between sample ranking within the tiers and disease trajectory. Fitted values correspond to the final pseudotime value.

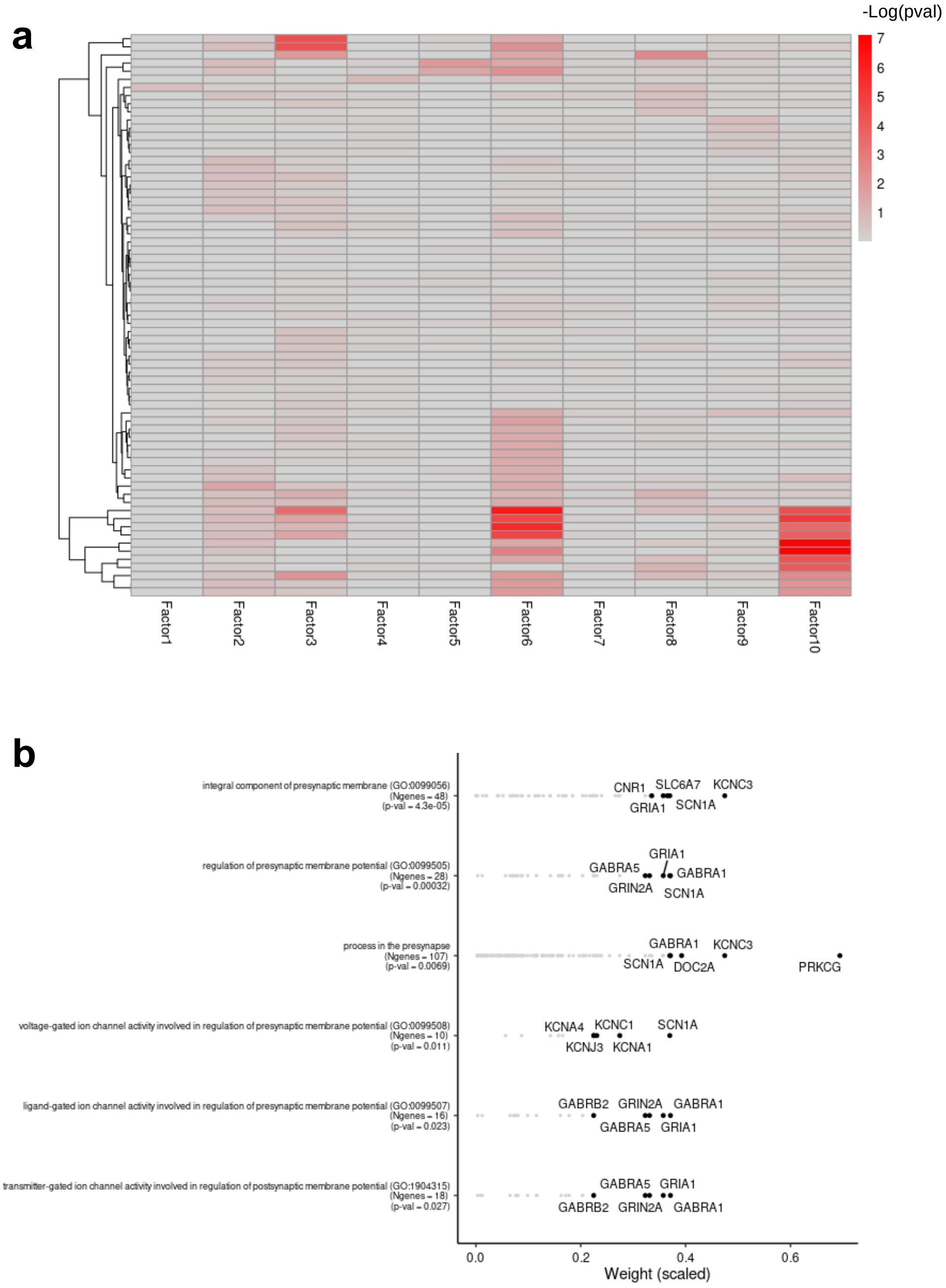


### Supplementary figure 8: MOFA factor SynGO enrichments.

**(a)** Heatmap with the adjusted -log(p-values) that result from the feature set enrichment analysis. Rows are SynGO[7] feature sets, and columns are factors. **(b)** Each row corresponds to a significant pathway from factor 3, sorted by statistical significance, and each dot corresponds to a protein. For each pathway, we display the top 5 proteins sorted by the weight in factor 3. The remaining proteins are coloured in grey.

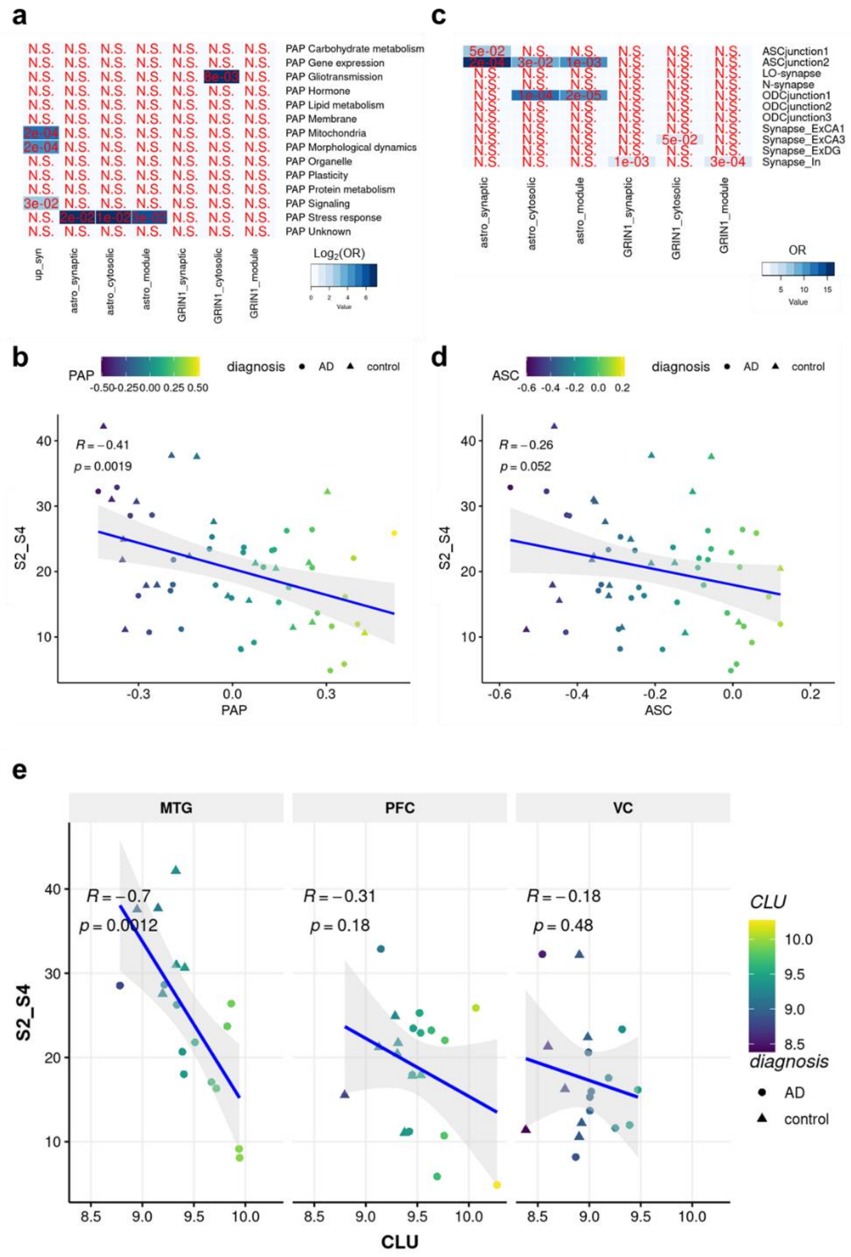


### Supplementary figure 9: PAP & ASC gene set activity.

**(a)** Heatmap of pairwise enrichment analysis of (upregulated proteins in synaptic fraction, ME2 astrocyte module, ME4 GluN1+ module) against PAP gene set[8]. Fisher's exact test determines the p-value and odds ratio compared to a genomic background (Colour= $\log_2$(OR), p-value). **(b)** Linear regression of the summed density of S2 and S4 synapse subtypes (S2_S4) against PAP activity (via GSVA) in the synaptic proteomics fraction. **(c)** Heatmap of pairwise enrichment analysis of (upregulated proteins in synaptic fraction, ME2 astrocyte module, ME4 GluN1+ module) against ASC gene set[9]. Fisher's exact test determines the p-value and odds ratio compared to a genomic background (Colour= OR, p-value). **(d)** Linear regression of the summed density of S2 and S4 synapse subtypes (S2_S4) against ASC activity (via GSVA) in the synaptic proteomics fraction. **(e)** Linear regression of the summed density of S2 and S4 synapse subtypes (S2_S4) against CLU abundance in the synaptic proteomics fraction.

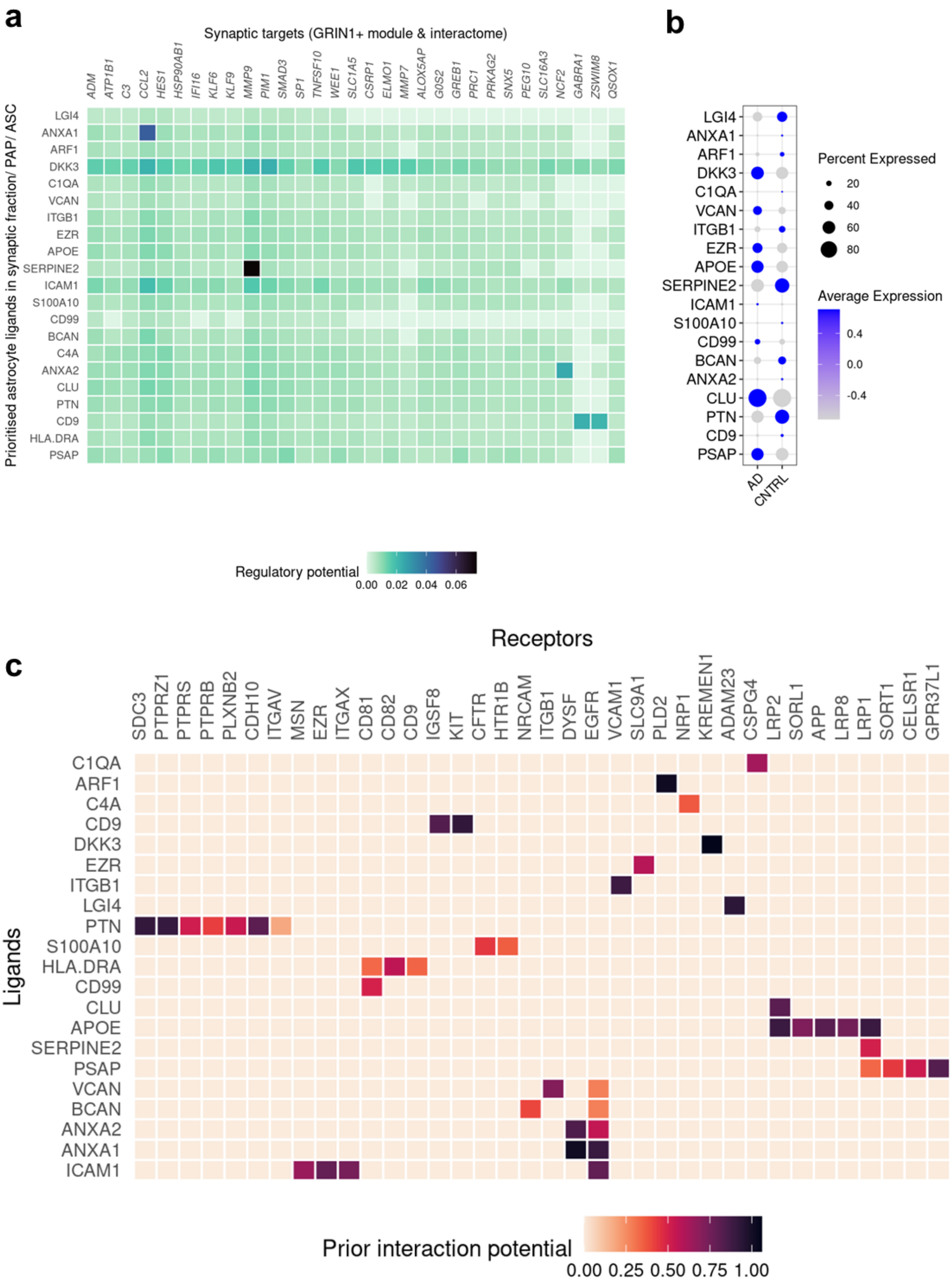


Supplementary figure 10: Astrocyte to synaptic targets communication.

**(a)** Heatmap showing NicheNet's ligand–target matrix denoting the regulatory potential between astrocyte/glia module ME2 / PAP / ASC ligands and target genes in the expanded GluN1[+] (GRIN1) module. **(b)** Average expression of prioritised astrocyte ligands in astrocyte single nuclei RNAseq data. Size indicates the percentage of cells expressing the genes, and colour indicates average scaled expression. **(c)** Heatmap showing NicheNet predicted upstream ligand-receptor pairs that might regulate genes in the extended GluN1+ module (module+interactome).

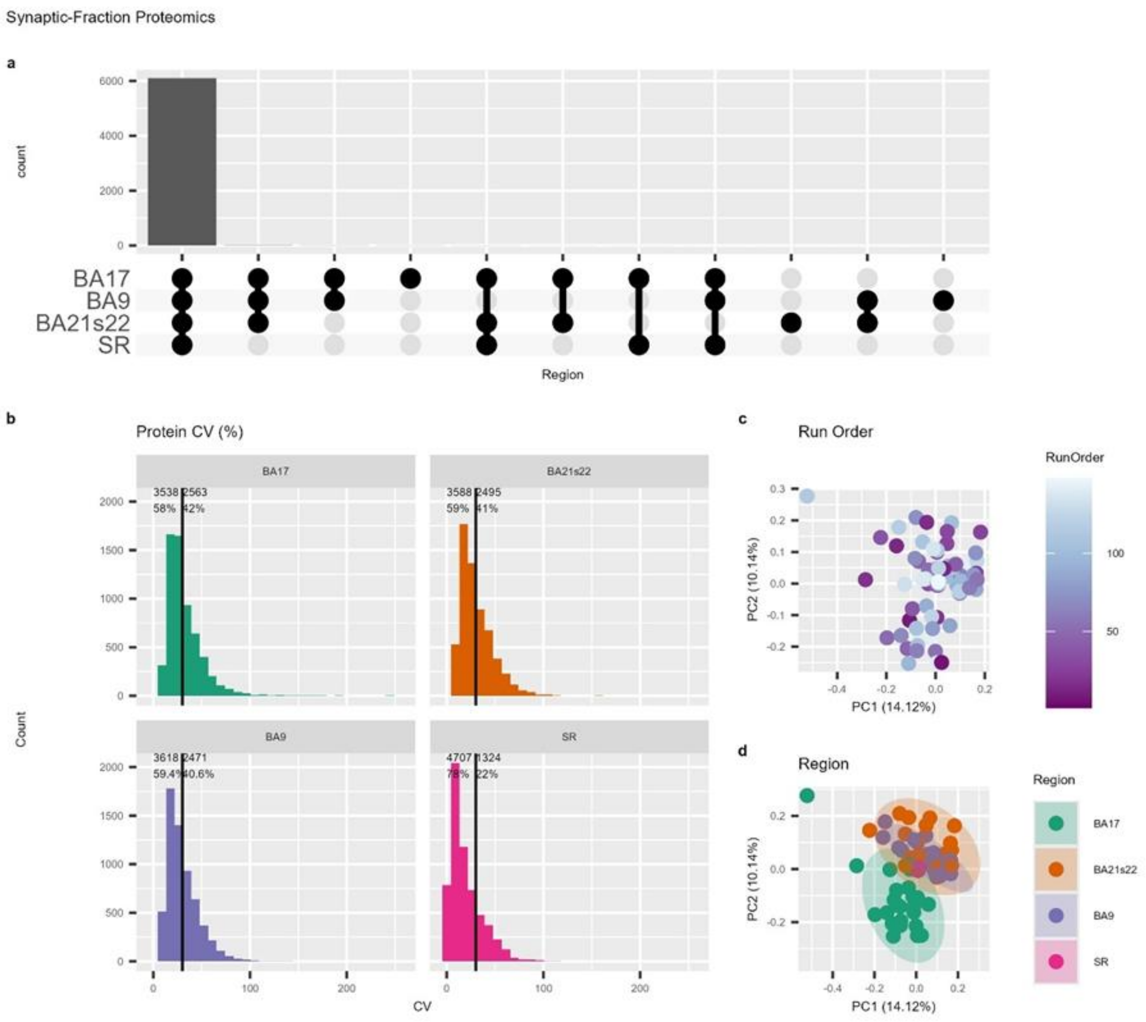


Supplementary figure 11: A summary of the synaptic proteomics data. **(a)** Upset plot showing proteins identified across brain regions (BA20s22, MTG; BA9, PFC; BA17, VC) and study reference (SR) **(b)** Protein coefficient of variation within distinct brain regions and SR. The number and proportion of proteins with CV < or > 30% is shown. **(c, d)** proteomics data annotated by **(c)** Run Order or **(d)** brain region.

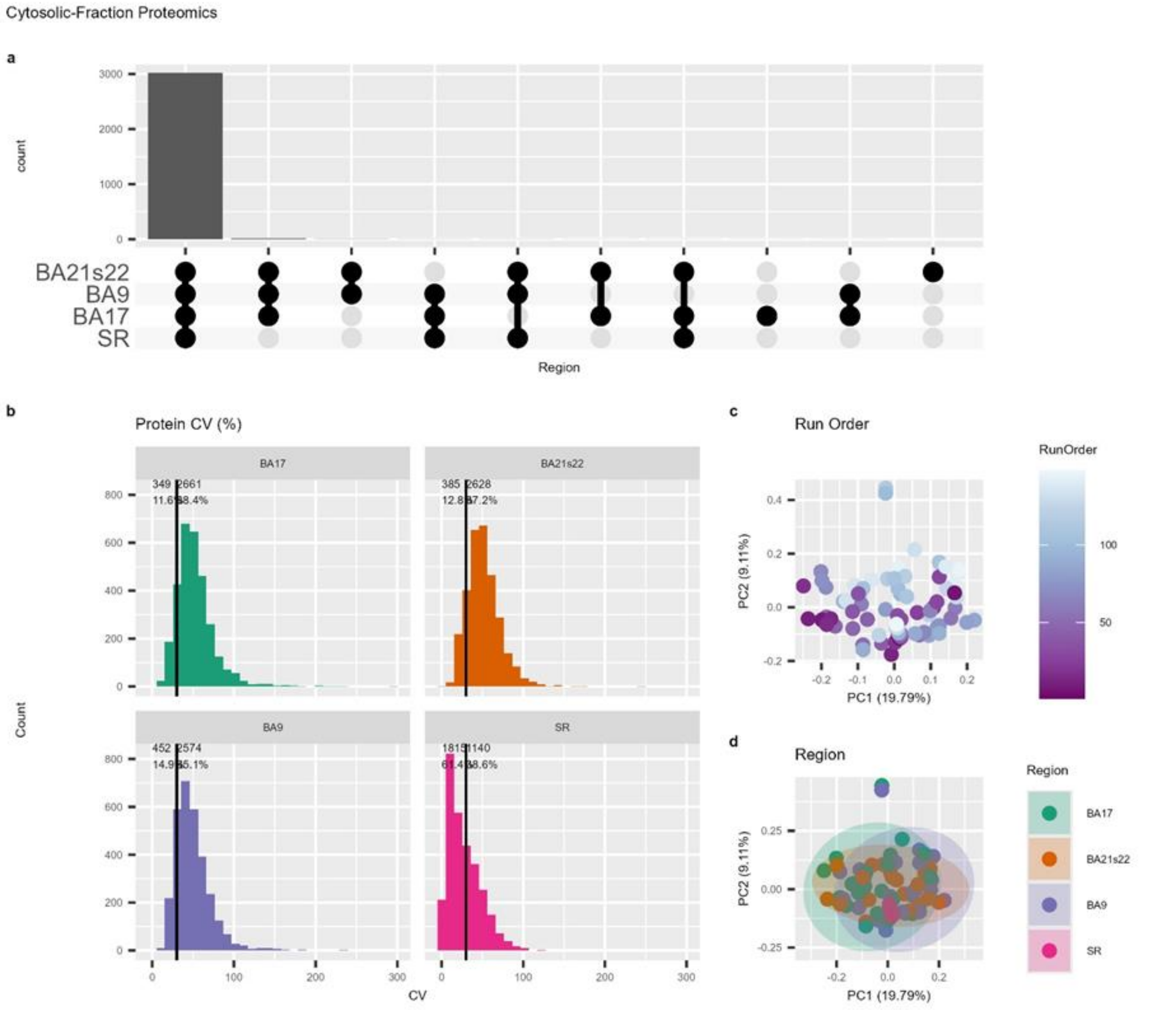


Supplementary figure 12: A summary of the cytosolic proteomics data. **(a)** Upset plot showing proteins identified across brain regions (BA20s22, MTG; BA9, PFC; BA17, VC) and study reference (SR) **(b)** Protein coefficient of variation within distinct brain regions and SR. The number and proportion of proteins with CV < or > 30% is shown. **(c, d)** proteomics data annotated by **(c)** Run Order or **(d)** brain region.

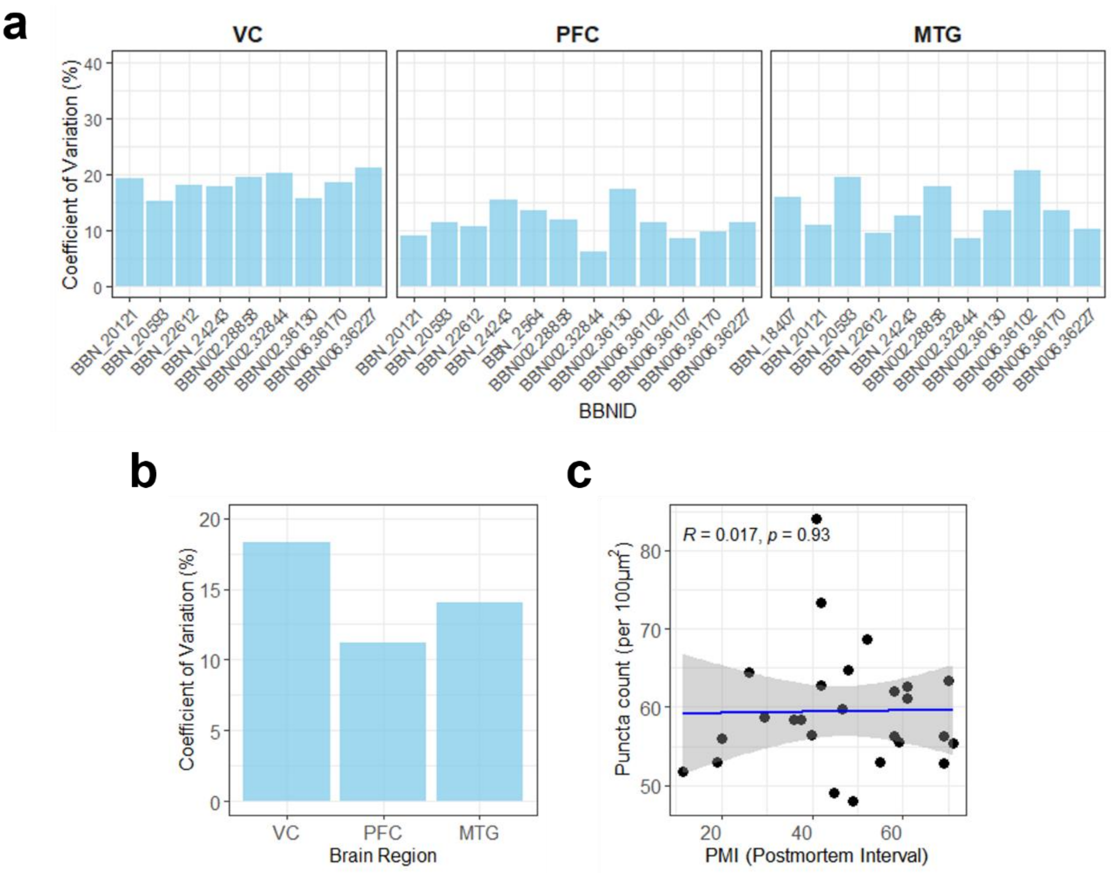


Supplementary figure 13: GluN1 synaptic puncta density is similar across the control cohort.
Bar plot displaying the percentage coefficient of variation (%CV) per region (VC, PFC, MTG) for GluN1+ synaptic puncta density across control samples used for synaptome mapping experiments. The relatively similar %CV values across cases (BBNID) within each region **(a)** and the less than 20% CV in each region **(b)** suggest that the variability is consistent and low. **(c)** Linear regression of the mean GluN1+ synaptic puncta density (per 100µm$^2$) against post-mortem interval (PMI). Pearson's correlation coefficient r value and p-value are displayed.
